# The lipidomic architecture of the mouse brain

**DOI:** 10.1101/2025.10.13.682018

**Authors:** Luca Fusar Bassini, Halima Hannah Schede, Laura Capolupo, Leila Haj Abdullah Alieh, Francesca Venturi, Alessandro Valente, Colas Droin, Daniel Trejo Banos, Irina Khven, Ece Z. Asirim, Anita Nashrallah, Irmak Kaysudu, Ekaterina Krymova, Giovanni D’Angelo, Gioele La Manno

## Abstract

Lipids are fundamental components of the brain, crucial for synaptic transmission and signal propagation. Altered brain lipid composition is associated with common and rare neuropathologies, yet, the spatial organization of the mammalian brain lipidome remains insufficiently characterized compared to other modalities^1–8^. Here, we mapped the membrane lipid architecture of the adult mouse brain at micrometric scale, across sexes, and during pregnancy. This Lipid Brain Atlas reveals that lipids define a fine-grained biochemical structure that aligns with functional anatomy. Membrane lipid spatial heterogeneity clusters into territories, which we termed *lipizones. Lipizones* partially mirror cell type territories, but also capture distal axon terminals. Through *lipizones*, (i) we reveal the organizing principles of the gray matter lipidome, related to connectivity and cytoarchitecture; (ii) we discover a new axis of oligodendrocyte heterogeneity in the white matter; (iii) and we find biochemical zonation in the choroid plexus and in the ventricular walls. We show that this lipidomic architecture can adapt to changing physiological needs. In the brain of pregnant females, the white matter is metabolically activated and the outer cortex is reorganized. These results are a foundational resource (https://lbae-v2.epfl.ch/), poised to reshape our understanding of lipids in brain development, physiology, and pathology.

## INTRODUCTION

Lipids account for the majority of the brain dry weight^9^. They form the cell membranes that constitute myelin, axons, dendrites, and synapses and enclose intracellular organelles and neurotransmitter-laden vesicles^10^. Seminal studies have examined the lipid composition of the mammalian brain across anatomical structures^11–14^. However, a systematic survey of the brain lipid metabolic architecture^15^ in relation to cell type composition, subcellular organization, functional anatomy, and connectivity is lacking.

To fill this gap, we used matrix-assisted laser desorption/ionization mass spectrometry imaging (MALDI-MSI)^16^, a technique capable of ionizing a micrometric portion of tissue and resolve for it a mass spectrum rich in diverse biomolecules^17^. We mapped the distribution of 172 lipids at quasi-cellular spatial resolution across 109 brain sections from 11 mice of 8 weeks of age, covering the entire brain volume. We identified 539 lipidome-defined brain clusters, which we termed *lipizones*. We characterized *lipizones* in relation to anatomy, cell type composition, cell compartments, connectivity, and biochemistry, revealing their biological organization and potential functional relevance. We investigated inter-individual and sex-related differences; finally, to explore how lipids vary in physiology, we charted the spatial lipidome in the brain of pregnant mice.

This study establishes the framework to investigate the lipid architecture of the healthy mouse brain and its changes in physiological settings, paving the way for future functional studies of brain lipids in evolution, development, pathology, and drug response.

## RESULTS

### A 3D lipidomic atlas of the mouse brain

We built a comprehensive lipidomic atlas of the adult mouse brain, measuring coronal sections from male and female brains. Using MALDI-MSI and the unified Mass Imaging Analyzer (uMAIA)^18^, we mapped 6,000 peaks across 7.2 million pixels (<5 µm laser spot at 25 µm raster interval) (***Figure 1a, b, Supplementary Figure 1a***; *see Methods*), which were warped into the Allen Brain Common Coordinate Framework (CCFv3^19^) for systematic comparison with anatomy and other modalities (***Supplementary Figure 1b-d***; *see Methods*). A total of 1,400 peaks passed noise quality control a combination of exact mass matching with liquid chromatography-tandem mass spectrometry (LC-MS/MS) and database searches^20,21^ (***Supplementary Figure 1e-j***; *see Methods*; ***Supplementary Tables 1, 2***).

**Figure 1:**
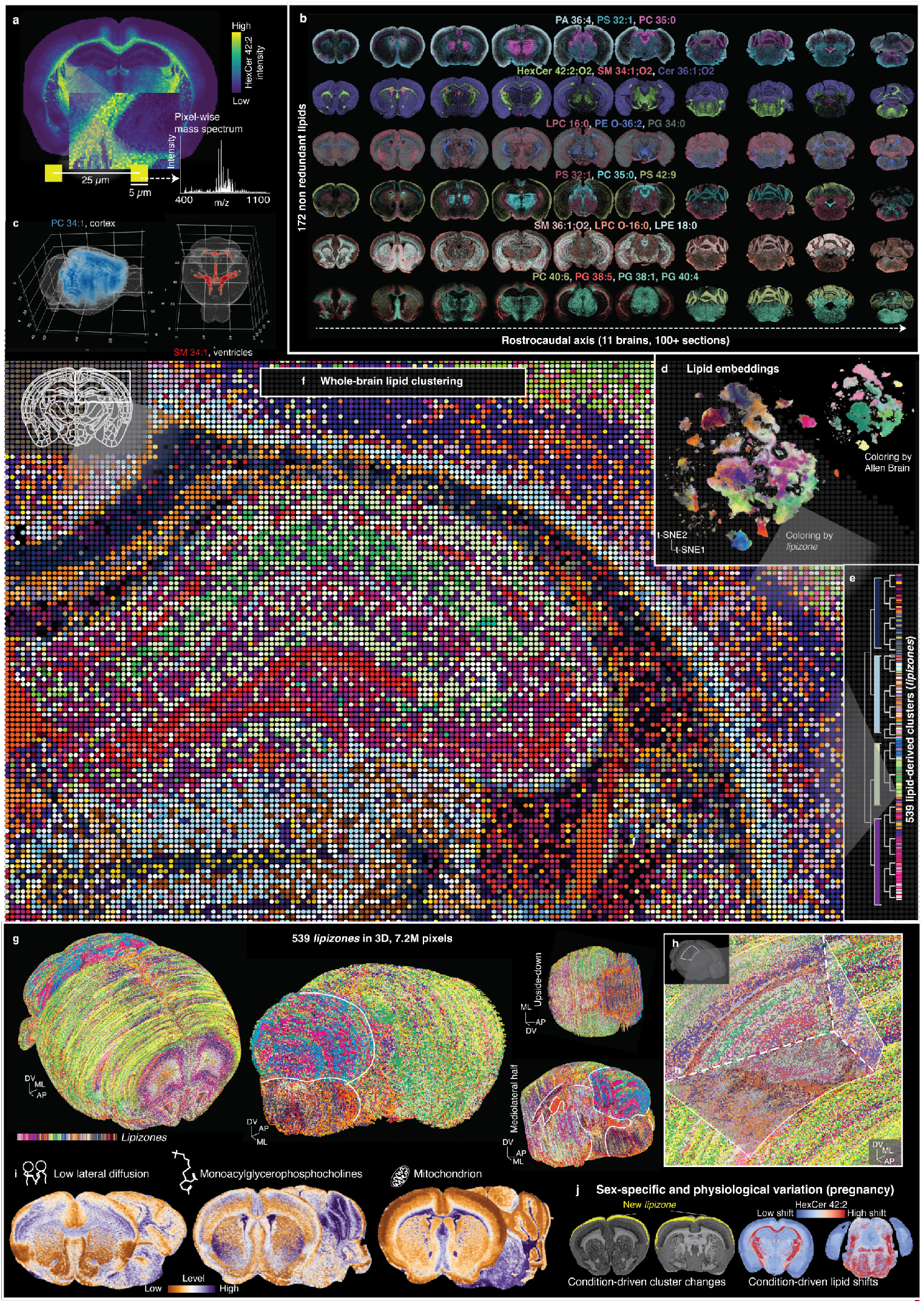
The mouse lipid brain atlas. (**a**) Schematic of a MALDI-MSI experiment measuring the mass spectrum of individual 5µm desorption points (pixels) along an adult brain coronal section; HexCer 42:2 distribution is displayed as an example. (**b**) Overview of MALDI-MSI composite images for selected annotated lipids along the rostrocaudal axis. (**c**) 3D distributions of selected lipids obtained interpolating serial sections. (**d**) t-distributed Stochastic Neighbor Embedding (t-SNE) of pixels in lipid space, colored by *lipizone* and by Allen Brain region. (**e**) Thumbnail of the (truncated) hierarchical tree of *lipizones*. (**f**) Zoom-in showing individual MALDI-MSI pixels as dots, colored by *lipizone*. Dots are enlarged for visualization purposes. (**g**) Whole-brain 3D visualizations of pixels colored by their corresponding *lipizone*. (**h**) Close-up revealing the *lipizone* structure near the hippocampal formation. (**i**) Spatial distributions for different lipid program scores estimated by the lipiMap algorithm. (**j**) Example of comparative results of lipidomic variations enabled by the atlas.

For most downstream analysis, we worked with 172 nonredundant lipids annotated with high confidence (*see Methods*). These included phosphatidylcholines (PCs: 31 species), phosphatidylethanolamines (PEs: 11), ether lipids (PE-Os: 10; PC-Os: 10), phosphatidylglycerols (PGs: 21), phosphatidylserines (PSs: 18), phosphatidylinositols (PIs: 14), phosphatidic acids (PAs: 12), lysophosphatidylcholines (LPCs: 10), lysophosphatidylethanolamines (LPEs: 6), sphingomyelins (SMs: 10), ceramides (Cers: 7), hexosylceramides (HexCers: 12), and other rarer classes (***Supplementary Table 2a***).

We first analyzed sections from two densely-sampled male brains (hereafter, ‘brain #1’ as a reference and ‘brain #2’ for validation). We then extended our analysis across additional sparsely sampled brains: three males from control, three females, and three pregnant (embryonic day E13.5) female brains. For pixel embedding and clustering, we used a set of lipids selected for high spatial variability, low dropout frequency, and consistency across tissue sections (***Supplementary Figure 1k, Supplementary Table 2b;*** *see Methods*). To capture the latent structure of the data, we applied non-negative matrix factorization (NMF), identifying 16 molecular signatures, followed by batch harmonization^22,23^ (***Supplementary Figure 2a;*** *see Methods*). Leveraging this latent space, we used eXtreme Gradient Boosting (XGBoost)-based imputation^24^ to recover lipid signals that were partially missing across sections (***Supplementary Figure 1l-o;*** *see Methods*). We then interpolated the data to reconstruct lipid distributions in three dimensions, resulting in a six-million-pixel 3D map covering 172 lipids (***Figure 1c***; *see Methods*). Subsequent Leiden clustering revealed spatially organized domains aligned with anatomical structures.

To capture finer substructures in the data, a binary hierarchical splitting algorithm was used to iteratively refine clusters based on both local and global lipidomic variability (***Figure 1d-h, Supplementary Figure 2b, c;*** *see Methods*). The process was stopped when branches lacked sufficient differential lipids or had too few pixels. The resulting clusters showed good agreement with Leiden clustering, but more robust to batch effects, and revealed finer substructure (***Supplementary Figure 2d-h***). We incorporated classification tasks (XGBoost) in the algorithm so that the clusters learnt on the first brain could be transferred onto the other brains (***Supplementary Figure 3a-d;*** *see Methods*); the transferred clusters were consistent with the clusters obtained reclustering independently brain #2 (***Supplementary Figure 3e***). Validated across 11 brains (biological replicates), our collection of methods is available in the Enhanced uMAIA for Clustering Lipizones, Imputation, and Differential analysis (EUCLID) package (see Methods; ***Figure 1a-j***). The Lipid Brain Atlas can be explored at https://lbae-v2.epfl.ch/ with the graphical user interface we built.

### *Lipizones* reveal a new axis of brain architecture

We identified 539 pixel clusters that we termed *lipizones* due to their remarkable spatial organization (***Supplementary Figure 4a-e, Supplementary Figure 5a, Supplementary Figure 6a-b, Supplementary Figure 7a***). The 539 terminal *lipizones* were hierarchically structured into 8 classes (splitting level 3), 31 subclasses (level 5), and 222 supertypes (level 8). *Lipizones* exhibited symmetry along the mediolateral axis, and all but two contained ≥74% of the 172 profiled lipids at detectable levels (***Supplementary Figure 4c***).

The primary division in the hierarchy separated gray and white matter-rich *lipizones* (***Figure 2a***). White matter *lipizones* localized to fiber tracts, hindbrain, and midbrain, as well as the thalamus and hypothalamus, and could be further divided into *lipizones* associated with oligodendrocyte-rich, neuron-poor regions and those linked to areas receiving extensive neuronal input.

**Figure 2:**
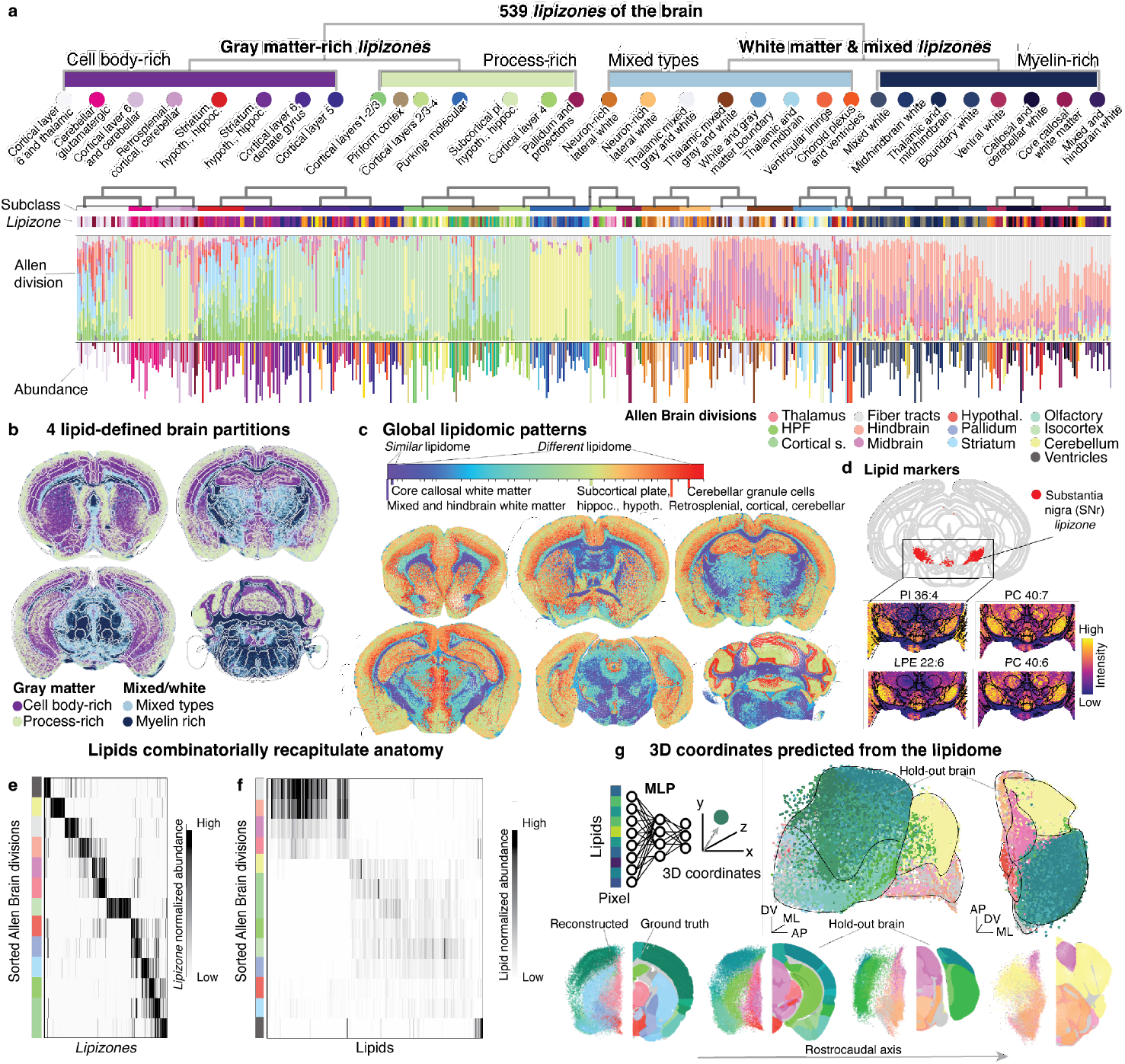
Comparison of *lipizones* and brain anatomy. (**a**) Summary visualization of the *lipizones* of the brain, including an annotated dendrogram of the 31 lipid-driven subclasses. For each *lipizone*, we display the distribution of its pixels across anatomical divisions and their abundance across the entire brain (highest values clipped). (**b**) Sections colored by the four major lipidomic subdivisions of the brain. (**c**) Sections with pixels colored by lipidomic similarity, i.e., similar color indicates similar lipidome. (**d**) Highlight on a *lipizone* that colocalizes with the *substantia nigra*, with the distribution of its local lipid markers. (**e**) Heatmap of *lipizone* normalized abundance by Allen division. (**f**) Heatmap of lipid normalized abundance by Allen division. (**g**) Summary of the analysis of pixel localization prediction based on lipidome. Top left: schematic of the approach. Top right: 3D visualization. Bottom: single section visualizations. Across all panels, pixels are colored according to their ground truth Allen Brain color.

Within the gray matter, *lipizones* were organized following an interior-exterior pattern (***Figure 2b***). The ‘interior’ group included cortical layers 5 and 6, granule cells of the cerebellum, most of the striatum, the majority of thalamic and hypothalamic nuclei, the periaqueductal gray, and parts of the hippocampus. The ‘exterior’ group encompassed cortical layers 1, 2/3, and 4, the piriform and entorhinal cortices, additional hippocampal regions, and the molecular layer of the cerebellar Purkinje cells (***Supplementary Figure 7b***; ***Supplementary Table 3***). The interior group predominantly comprised anatomical regions enriched in glutamatergic neurons, whereas the exterior group included areas with abundant GABAergic populations.

Intriguingly, the sequential formation of layers during brain development^25^ may explain the early bifurcation between deep and mid-surface cortical layers. However, the reason for lipidomic similarity between cerebellar Purkinje arborization and outer cortex, as well as between granule cells and inner cortex, remained unexplained.

This subdivision was driven by myelin lipids (HexCers and Cers), more abundant near fiber tracts and *arbor vitae*, i.e., in the inner cortical layers and the granule cells of the cerebellum. The relative enrichment of excitatory synaptic membranes^26^, particularly in areas with few cell bodies (***Supplementary Figure 7c***), such as cortical layer 1 and the Purkinje molecular layer^27^ may contribute to this major structural division.

As the hierarchy displayed structured anatomical organization across the entire brain (***Figure 2c***), we named the *lipizones* after the Allen Brain regions in which they were most enriched, for example, ‘DG-po-DG-sg’ (dentate gyrus polymorphic layer and subgranular zone) and ‘ICd’ (dorsal inferior colliculus) (***Supplementary Figure 3f;*** *see Methods*). Most *lipizones* encompassed between 800 and 25000 pixels across the entire atlas (***Supplementary figure 3g***). *Lipizones* showed a strong bidirectional association with the 13 major divisions of the Allen Brain Atlas (***Figure 2e; Supplementary Figure 4d***), a pattern only partially exposed by individual lipids (***Figure 2f***).

Exceptions were PE-O 38:7, restricted to the isocortex; SM 36:2, localized to the amygdala and the hippocampus; SM 34:1 and SM 38:1, confined to the ventricles; PC 40:6, enriched in the cerebellum (***Supplementary Figure 7d***). Interestingly, PC 40:6, already known to be enriched in the cerebellum^28^, is the major lipid containing docosahexaenoic acid (DHA), and it is decreased in aging and Acyl-CoA Synthetase Long Chain Family Member 6 (*Acsl6*)–/– mice, with cerebellar distress and hyperlocomotion^28^. The scarcity of single-lipid markers likely reflects the fact that distinct metabolic states modulate the relative abundance of membrane lipids, rather than creating all-or-nothing patterns of individual lipids.

Hierarchical clustering revealed splits of varying lipidomic divergence. The contribution of individual lipids varied across the hierarchy: some showed differential expression across many splits, while others appeared only at deeper levels (***Supplementary Figure 2c***). Notably, fold changes rarely exceeded twofold, reinforcing the idea that lipid-based partitions are driven by combinatorial patterns rather than single-lipid dominance.

When examining inter-individual variation in the three sparsely sampled males and three females, we found that for 90% of the lipids, only 1.1% to 6.3% of the total subclass variance could be attributed to individual differences (*see Methods*). The most variable lipids included LPE 22:6, LPE 22:4, and PC 40:1, while the most stable were PE-O 36:1, LPE 20:0, and Cer 40:1 (***Supplementary Figure 7e***). This minimal inter-individual variation reflects the fact that the analyzed lipids are stable structural components of cellular membranes, tightly regulated to preserve membrane integrity and function.

The association between *lipizones* and brain anatomy extended to the 616 fine-grained regions of the Allen Brain Atlas (***Supplementary Figure 7f***). Nevertheless, exceptions emerged: some *lipizones* displayed spatially coherent distributions that crossed conventional anatomical boundaries (***Supplementary Figure 7g***). *Lipizones* often recapitulated specific tissue niches, for example, the *substantia nigra reticulata*, distinctly marked by several phospholipids, including LPE 22:6, PI 36:4, PC 38:7, and, importantly, PC 40:6 (***Figure 2d***). Hippocampus and cortical subplate, which derive developmentally from the pallium and are transcriptionally related^2^, shared several *lipizones* (***Figure 2e***). Similarly, pallidum and striatum, both derived from medial/lateral ganglionic eminences of the subpallium^29^, shared many *lipizones*.

Having established the presence of structured regional heterogeneity, we next asked whether lipid distributions encode enough information to predict anatomical location. To test this, we trained a multi-layer perceptron on brain #1 to predict 3D spatial coordinates using only lipid profiles as input (***Figure 2g, Supplementary Figure 7h;*** *see Methods*). The model generalized well to the entirely held-out brain #2, reconstructing regional boundaries without relying on spatial information. Collectively, these results show that brain lipid composition carries spatially organized, fine-grained information, and that *lipizones* constitute biologically meaningful units.

### Brain lipids are irreducible to the transcriptome

We next wondered whether the spatial heterogeneity of the lipidome is directly related to the expression patterns of metabolic enzymes. We integrated the Yao et al. (***Supplementary Figure 8a***) and the Langlieb et al.^1,3^ spatially-resolved brain cell atlases with our lipid atlas based on the CCF registration. After filtering well-matched sections and cells, we investigated whether enzymes and the membrane lipids they produce are correlated at the cell type level. We extracted a reagent-product-enzyme list for the lipids in this study using LINEX2^30^ followed by expert curation (*see Methods*), and we used enzyme transcripts from scRNA-seq as proxies for enzymes. Except for 81/338 pairs of an enzyme and its product correlating R > 0.5, enzyme transcripts were not trivially positively correlated (R = −0.13 ± 0.57) with the lipid products of the reactions they catalyze (***Figure 3a***). On the other hand, enzyme expression profiles might help to disambiguate isomers that are indistinguishable by MALDI-MSI. This was the case for the HexCers galactosyl- and glucosylceramide, synthesized by Ugt8a and Ugcg, respectively. We found that Ugt8a expression was highly correlated with HexCers, suggesting that galactosylceramide comprises a major portion of adult brain HexCers.^31^

**Figure 3:**
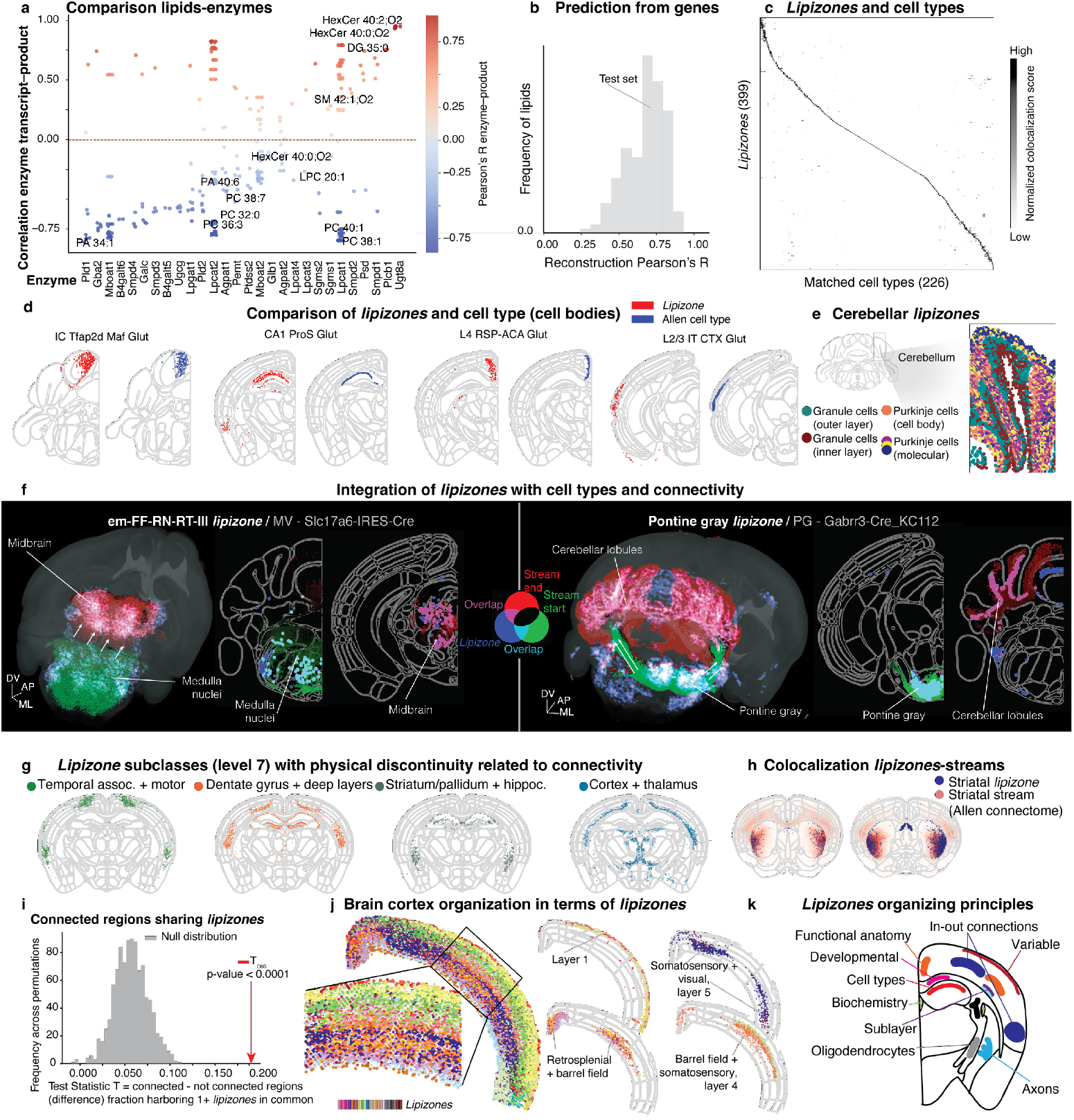
Organizing principles of the brain lipidome. (**a**) Strip plot reporting the Pearson’s correlation between enzyme expression and corresponding lipid products (multiple products per enzyme). (**b**) Histogram of reconstruction accuracies (Pearson’s R) of lipid levels from the transcriptome via XGBoost regression. (**c**) Colocalization matrix (reciprocal enrichment) for matching cell type territories and *lipizones*. (**d**) Spatial plots of *lipizones* and the cell type territories they colocalize with. (**e**) Spatial zoom-in on the granule cells layer of the cerebellum and its neighborhood, colored by *lipizones* named based on their locations. (**f**) 3D and 2D visualizations of two representative *lipizones* that likely include cell bodies (cyan) and their distal terminals (magenta). The associated connectomic streams are represented in background, colored from green to red, passing through black. The pixels of the *lipizones* that do not overlap with the connectomic streams are colored in blue. The gray text in the titles contains the connectome experiment injecton site and mouse line. (**g**) Spatial plots of selected *lipizone* subclasses (level 7) spanning anatomically distant yet connected brain regions. (**h**) Spatial plot showing the overlap between a striatal *lipizone* and a connectomic stream. (**i**) Histogram of the bootstrap distribution for the frequency of sharing a *lipizone* for two Allen anatomical regions. Observed value for regions with high connectivity displayed with a red arrow. (**j**) Spatial plot of cortical *lipizones*. Left: close-up. Right: examples, displayed in groups spanning the same anatomical regions. (**k**) Schematic illustrating the lipidome organizing principles.

Our results show that lipid distributions cannot be reliably inferred by individual enzyme expression. Reasoning that a more holistic view of the metabolic network might improve predictions, we tested whether supervised models could leverage covariation between metabolic genes and lipid distributions. We used canonical correlation analysis (CCA) and redundancy analysis with the Allen Brain MERFISH-imputed 500+ metabolism-related genes^1^, finding that metabolic genes linearly captured ∼36.6% of the total variance in lipid profiles (***Supplementary Figure 9a***). Finally, to comprehensively assess the information content relationship between gene expression and lipidome, we quantified the predictability of lipid distributions from the Allen Brain MERFISH-imputed full transcriptome^1^ using XGBoost regression to capture nonlinear relationships (*see Methods*). Broad lipid distributions (R^2^ > 0.25) could be predicted for 86.7% of lipids, with 35/172 lipids being predicted with good accuracy (R^2^ > 0.60, ***Figure 3b***). Considering a conservative upper bound on prediction performance of R^2^≈0.81 based on empirically estimated irreducible noise in lipid-to-lipid prediction, the gene-based model (test-set average R=0.68) captured 56% of the explainable lipid variance (*see Methods*). These results suggest that the brain lipidome is irreducible to the transcriptome. The membrane lipidome likely reflects cell states happening on time scales only partially overlapping with those of gene expression.

Feature importance analysis (*see Methods*) across 251 anatomical regions and 172 lipids revealed that predictive genes were significantly enriched for cell type markers, with 91% region-lipid pairs significantly enriched for markers (bootstrap, BH-FDR < 0.05, ***Supplementary Figure 9b***). For example, *Th* had high feature importance in the *substantia nigra, Drd1* in the striatum, *Slc6a1* in cortical layer 4, *Prox1* in the hippocampus, *Pcp2* in the cerebellum, and *Gad2* in the thalamus (***Supplementary Figure 9c***). This indicates that when employing complex nonlinear functions across the entire transcriptome, reliable, yet imperfect, lipid predictions derive from the model focusing on cell type identity, suggesting that lipid distributions reflect properties intrinsic to cellular lineages rather than transient states. These could include shared developmental history, non-transcriptional regulatory mechanisms, metabolite accumulation, structural organization, and network connectivity patterns.

### *Lipizones* capture the bodies and terminals of cell types

MALDI-MSI pixels contain a mixture of cell bodies, axons, dendrites, extracellular matrix, and glial projections, all of which may in principle contribute to *lipizones*. First, to test how *lipizones* corresponded to cell type territories in terms of cell bodies, we measured colocalization at all hierarchical levels by reciprocal enrichment analysis (*see Methods*). We found a striking correspondence (***Figure 3c***): excluding the 65 fiber tract and the 57 cerebellar *lipizones*, 76% (304/399) *lipizones* discovered by our clustering algorithm overlapped with cell type territories. Interestingly, cell types such as “3283 IC Tfap2d Maf Glut_1”, “0349 L4 RSP-ACA Glut”, “0116 L2/3 IT CTX Glut_3”, and “0262 CA1-ProS Glut_1” were rediscovered by our lipid-only clustering method (***Figure 3d; Supplementary Figure 9d***). By exploiting the correspondence between *lipizones* and cell type territories, we were able to estimate the lipidome of these cell types (***Supplementary Figure 9e***).

66 cell type territories were matched by more than one *lipizone*. Given their spatial coherence, these *lipizones* might represent different subcellular regions, myelination levels, or “lipotypes”^32^. One such example is the territory of the granule cells layer of the cerebellum (***Figure 3e***). The inner sublayer (proximal to the *arbor vitae*), where myelinated mossy fibers form synaptic glomeruli, was enriched in myelin-related lipids^33^, while the outer layer was relatively enriched in phospholipids (***Supplementary Figure 9i***). The *lipizone* hierarchy grouped the inner granule cells layer with regions rich in cell bodies including cortical layer 6. In contrast, the outer granule cells layer, clustered with axon-rich regions including striatal, hypothalamic, and thalamic areas, suggesting that the double-layer might reflect compositional differences between neuronal cell bodies and axonal projections^34^. The *lipizones* that did not overlap with cell type territories were largely localized to the hindbrain, striatum, and midbrain (regions that are rich in myelin), and hypothalamus, a region covered for 33% by only two *lipizones*.

*Lipizones* that recapitulated cell type territories often extended to physically distant regions, suggesting that individual *lipizones* might capture cell bodies and their distal axon terminals. We therefore sub-clustered individual *lipizones* based on spatial coordinates, and used the Allen cell locations, corresponding to cell bodies, to classify subclusters as cell body- or terminal-related (*see Methods*). To support the role for soma- or terminal-*lipizone* subclusters, we further examined the Allen connectomic streams^4^ that overlapped significantly with both subclusters, checking whether the subclusters were localized at opposite extremes of at least one stream. Based on these data, for 106 *lipizones*, we found initial evidence of two subclusters defined by physical space, directionally connected, with one subcluster colocalizing with the cell bodies (***Figure 3f; Supplementary Figure 9f***). Differently from cell body-based spatial transcriptomics, lipidomic profiles are therefore able to capture long-range projections within individual cell types.

Notably, regions with known functional connections often clustered together in our data-driven hierarchy despite their spatial separation (***Figure 3g***). For example, ectorhinal, temporal association, and motor cortices, all forming predominantly corticocortical connections, shared similar *lipizones*^*4*^. Similarly, the hippocampus clustered with its known input and output regions, including lateral septal nucleus, cortical amygdala, entorhinal cortex, and cortical motor areas^35–37^. Anterior cingulate and hypothalamic nuclei^38^ have similar lipidomes, as well as thalamic nuclei and cortical layer 6, which form established corticothalamic circuits^39,40^. Although thin individual axons (overwhelmed by the myelin wrapping around them) are probably not yet within reach as well-resolved entities, we also found spatial correspondence between some specific *lipizone* distributions and axon streams, such as in the striatum (***Figure 3h***).

Compelled by these observations, we compared the mesoscale brain connectivity^4^ with *lipizones* (structured permutation test of the connectivity density of brain regions sharing *lipizones* against random regions; *see Methods*). We found that connected brain regions shared one or more *lipizones* more frequently than by chance (p < 0.0001) (***Figure 3i***). Collectively, these findings suggest that the organization of *lipizones* captures fundamental aspects of the brain connectivity architecture.

In the cortex, we identified spatial distributions of *lipizones* that recapitulated cortical layering while revealing organizational patterns not observed in the Allen cell type atlas^1^ (***Figure 3j, Supplementary Figure 9g***). Notably, we discovered 14 layer 5 barrel field *lipizones* that were not detected by gene expression (***Figure 3j***). In layer 5, these *lipizones* were also enriched in the retrosplenial cortex, a region that was proposed to connect with the barrel cortex.^41^ The barrel field and the retrosplenial cortex were jointly enriched in specific lipids, such as Cer 42:2 and SM 42:3 (***Supplementary Figure 9h***). We also identified *lipizones* restricted to the barrel field and somatosensory cortex in layer 4, while other *lipizones* spanned physically distant regions, such as *lipizones* crossing simultaneously the somatosensory and visual cortices. These findings suggest that lipid distributions might capture functional specialization across cortical areas that extends beyond the cell type composition captured by the current transcriptomics methods.

In conclusion, *lipizones* model biomolecular variability that correlates with cell type composition, cytoarchitecture, and circuitry (***Figure 3k***), revealing the spatial patterns of cell bodies and their terminals. This makes *lipizones* valuable, non-redundant candidate units for examining the metabolic and structural architecture of the brain, as future functional studies will ultimately establish.

### Structured diversity of myelinated regions

We hypothesized that *lipizones* could reveal unrecognized heterogeneity in regions with sparse cell bodies such as highly myelinated areas. Comprehensively, a branch of the hierarchy encompassing 65 *lipizones* localized to fiber tract regions, *arbor vitae*, and the hindbrain. These regions were the richest in myelin, as confirmed using two marker scores—hexosylceramide and oligodendrocyte membrane lipid score (derived from a previous study^11^; see Methods, ***Supplementary Figure 10a***). Individual *lipizones* were specifically restricted, for instance, to the periventricular white matter, fimbria, dorsal callosum, *arbor vitae*, internal capsule, optic tract, corticospinal tract, and anterolateral callosum (***Figure 4a***; ***Supplementary Figure 10b, c***).

**Figure 4:**
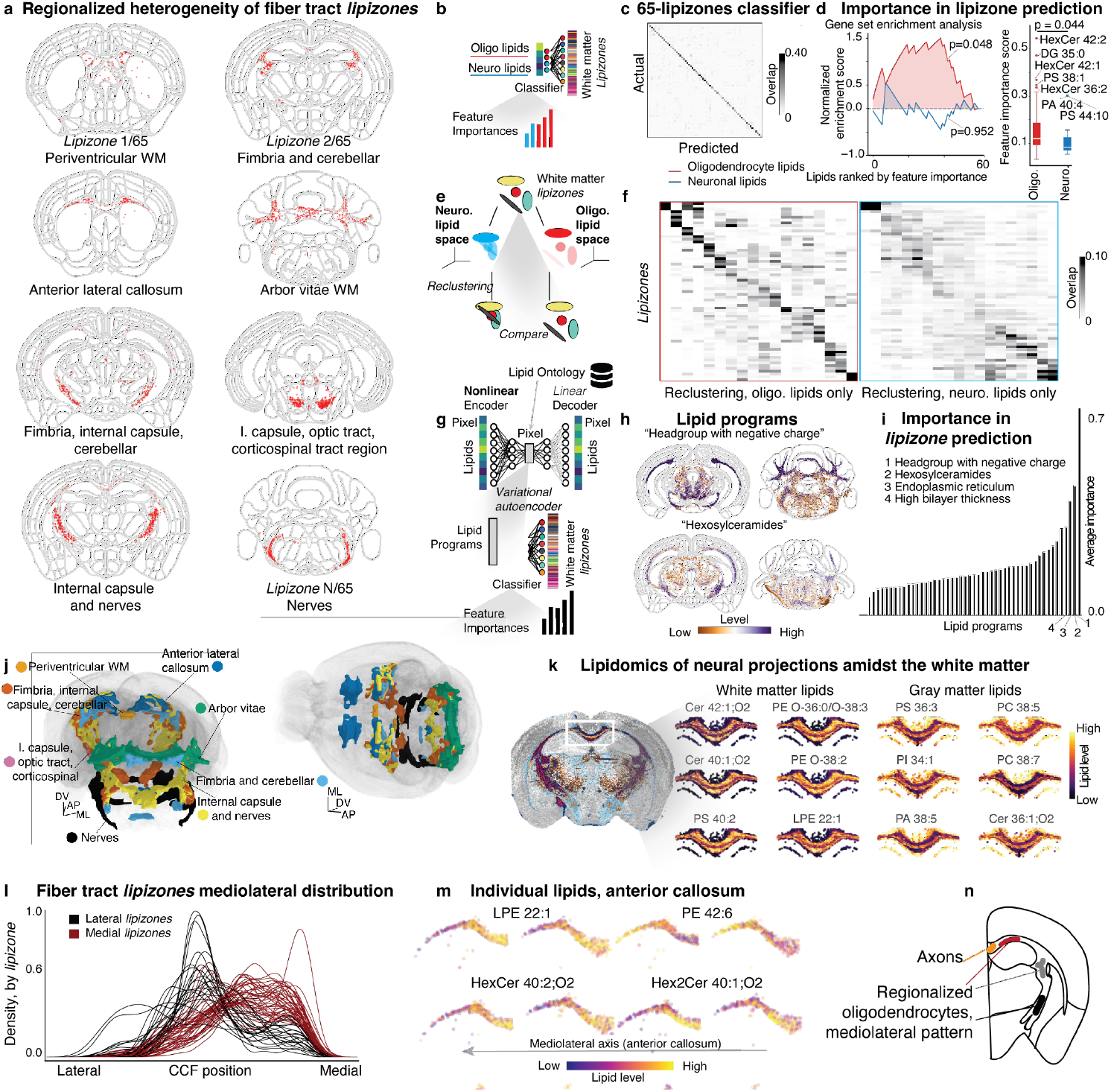
White matter lipidomic heterogeneity. (**a**) Spatial plots of eight distinct regionalized white matter *lipizones*. (**b**) Schematic of the *lipizone* classification task to extract the relative importances of neuronal and oligodendrocyte lipids in determining white matter heterogeneity. (**c**) Heatmap for the normalized confusion matrix ground truth vs predicted for the 65-class *lipizone* classification task, evaluated on a held-out set. (**d**) Gene set enrichment analysis (GSEA) plot showing the ranks of oligodendrocyte and neuronal lipids in the important features set for the 65-class *lipizone* classification task. The boxplot on the right shows the distributions and outliers for the feature importances of the two sets, with the top lipid predictors highlighted. (**e**) Schematic of the pixel reclustering task using neuronal and oligodendrocyte lipids only as alternative feature sets. (**f**) Heatmaps of the normalized confusion matrices comparing the 65 *lipizones* with the clusters computed from the feature sets of oligodendrocyte-specific lipids (left) and neuron-specific lipids (right). (**g**) Top, schematic of the biochemically-constrained variational autoencoder (lipiMap), which, using the Lipid Ontology database, learns latent biological programs from the lipidome. Bottom, schematic of the *lipizone* classification task to extract the relative feature importances of lipid programs in determining white matter heterogeneity. (**h**) Spatial plots of two lipid programs with regional heterogeneity within the white matter. (**i**) Barplot of the feature importances of lipid programs for the white matter *lipizone* classification task, highlighting the most important programs. (**j**) 3D renderings of eight white matter-related *lipizones*, with color-coding. (**k**) Spatial plot highlighting the *lipizones* forming a bundle passing through the fiber tracts in the anterior callosum, with zoom-ins colored by differential lipids between the bundle and its surroundings. (**l**) Kernel density estimates of white matter-related *lipizone* abundances along the mediolateral axis, one curve per *lipizone*, with color coding highlighting medial and lateral *lipizones*. (**m**) Zoom-in spatial plots of lipids with a mediolateral gradient in the anterior callosum (views are limited to one hemisphere). (**n**) Schematic representing the discoveries related to the white matter.

This regionally-structured diversity of *lipizones* is remarkable, particularly given that myelin has been described as homogeneous in previous reports and fewer than ten oligodendrocyte subtypes are recognized transcriptomically^1,42^. We therefore asked whether this heterogeneity depends on myelin or axon composition. To do so, we trained a classifier using as features only oligodendrocyte- and neuron-enriched lipids^11^. We then used SHAP (SHapley Additive exPlanations)^43^ to extract the feature importances for the prediction of each *lipizone* (***Figure 4b;*** *see Methods*). We first confirmed that the 65 white matter *lipizones* can be distinguished from one another by training a 65-class classifier which was able to recover them in this all-vs-all setting (***Figure 4c***). Looking at the feature importance set, oligodendrocyte-restricted lipids dominated compared to neuronal-restricted lipids (gene set enrichment analysis, p=0.048 for oligodendrocyte lipids, non-significant for neuronal lipids; t-test, p=0.044) (***Figure 4d***). As a confirmation, we limited the feature space to oligodendrocyte or neuronal lipids only, and we reclustered by principal component analysis (with 10 components for both feature spaces) and Leiden algorithm (***Figure 4e***). We found that the oligodendrocyte lipids space was better at retrieving the fine-grained *lipizones* (Normalized Mutual Information = 0.24 vs 0.12; Adjusted Rand Index = 0.057 vs 0.023) (***Figure 4f***).

To reveal the biological processes underpinning the observed heterogeneity, we designed lipiMap, a biochemically-constrained variational autoencoder (***Figure 4g;*** *see Methods*). This architecture was inspired by expiMap^44^, a biologically-informed deep learning framework, which we extended to obtain a latent representation constrained by known lipid ontologies^45^. We trained the model on brain #1, enriching each pixel with interpretable lipid programs^44^ (***Figure 4h; Supplementary Figure 11a-c***). We were able to study biophysical and biochemical properties in space, including lipid chemical structure, organelles, signaling, lateral diffusion, membrane transition temperature, and thickness. We inferred thicker membrane properties in the white matter, accompanied by a low lateral diffusion, aligning with the notion that white matter membranes are stable and compact (***Supplementary Figure 11d***). We found that known myelin-related characteristics had the highest feature importance to distinguish the white matter-rich *lipizones*, including the programs “headgroup with negative charge”, “hexosylceramides”, and “high bilayer thickness”^46^ (***Figure 4i***). Collectively, these observations suggest that the lipidome of oligodendrocytes drives the regional heterogeneity of white matter-rich *lipizones* (***Figure 4j***).

We next asked whether axonal lipid composition contributes to residual heterogeneity in fiber tract *lipizones* outside the major myelin-rich branch. We focused on a specific part of the white matter that carries interhemispheric axons within the corpus callosum^47^. We found that *lipizones* in this area identified a bundle that crosses between the two hemispheres (***Figure 4k***). In fact, differential analysis on local *lipizones* partitioned the bundle, rich in neuronal marker Cer 36:1^11^ and phospholipids, from the surrounding white matter, rich in myelin lipids including other ceramides, PSs and ether lipids (***Figure 4k, Supplementary Figure 10d***). This pattern with lightly myelinated axons in the middle, previously reported for the GD1a and GT1b axonal gangliosides^48^, suggests that variable degrees of myelination are a source of heterogeneity.

Given the stability of myelin and its retention of ontogenic marks^49–51^, we hypothesized that white matter *lipizone* diversity might reflect developmental origin. Since myelination extends into early adulthood, we reasoned that its lateral-to-medial anterior developmental waves in the callosum might result in enduring signatures in the adult^52,53^. Interestingly, we found that anterior fiber tract *lipizones* largely grouped into lateral and medial pools (***Figure 4l; Supplementary Figure 10e***) (average CCF 3.72 vs 4.33, t-test p<0.0001). Specific lipids, including HexCer 40:1 and 40:2, LPE 22:1, and PE 42:6, correlated with the mediolateral axis in the anterior callosum (***Figure 4m***), suggesting, at a correlative level, that myelin lipid signatures might retain chronotopic developmental features (***Figure 4n***).

### The ventricular system is metabolically zonated

In the ventricles, metabolism supports immune homeostasis and detoxification^54^, cerebrospinal fluid (CSF) is produced and filtered^55^, and adult neurogenesis takes place^56^. Since these regions harboured a high residual variability unexplained by our terminal *lipizones* (***Figure 5a***), we further clustered the ventricular and periventricular *lipizones*. The clustering resulted in 15 subzones, including white matter-related ventricular portions (likely, acellular ventricular fillings), which we annotated based on their spatial distribution (***Figure 5b-d***). We found *lipizones* reminiscent of vascular leptomeningeal cells (VLMCs), ependymal cells, and telencephalon astrocytes spatial organization. These ventricular *lipizones* were characterized by different lipid families, with TGs enriched in the white matter-related ventricular filling, and phospholipids and lysophospholipids in the choroid plexus (***Figure 5c***).

**Figure 5:**
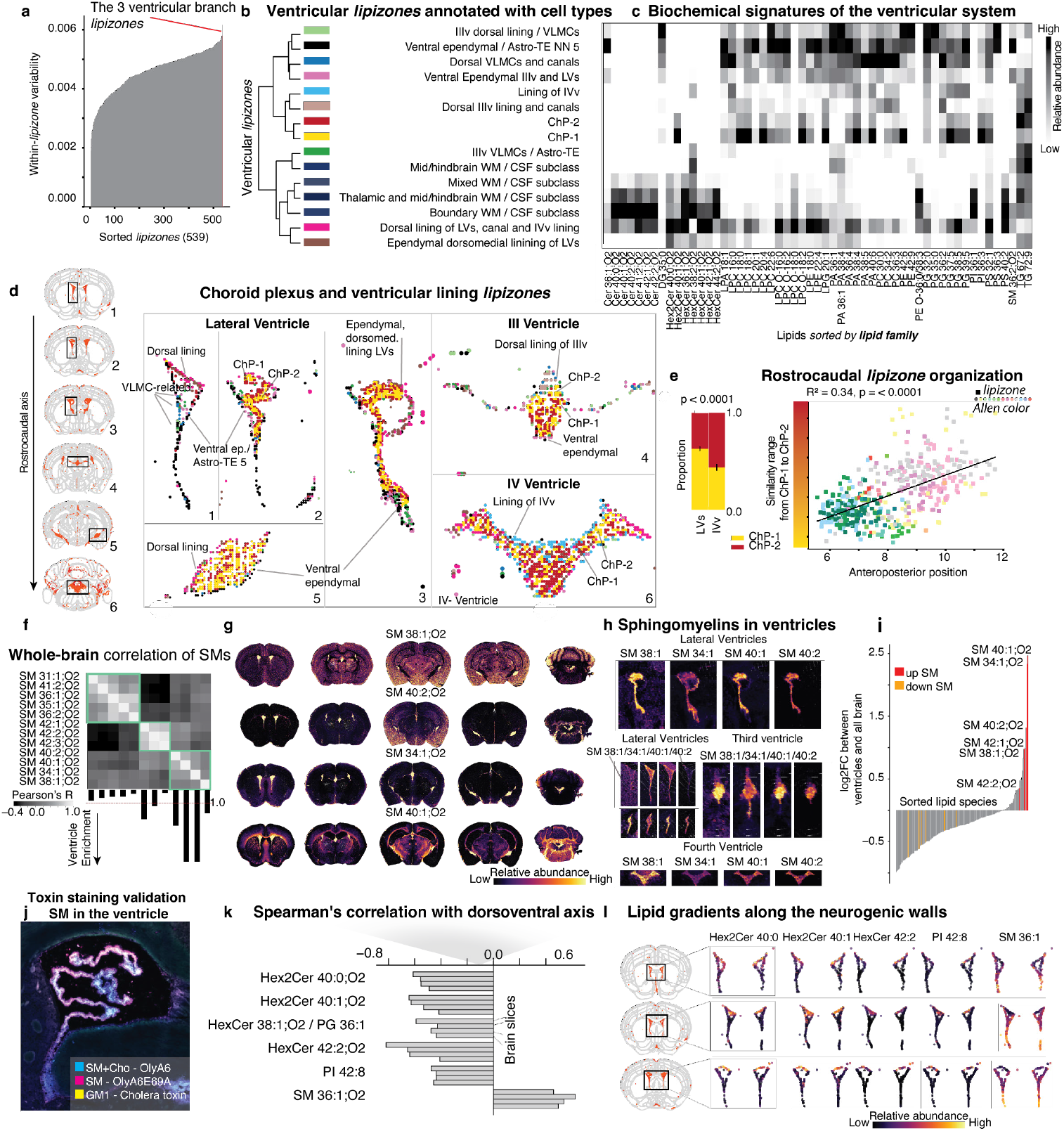
Metabolic zonation in the ventricular system. (**a**) Lineplot of within-*lipizone* variability for all the 539 *lipizones*. Lines are sorted in ascending order. Venticular *lipizones* are indicated in red. (**b**) Annotated dendrogram representing the hierarchy of fine ventricular *lipizones*, with (**c**) matched heatmap of lipid relative abundances. Lipids are ordered by lipid class. (**d**) Close up on ventricles, and their *lipizones*. Lipizones are colored as in (b). (**e**) Left: stacked barplots of the proportions of the two core choroid plexus *lipizones*, in the lateral (LVs) and fourth (IVv) ventricles. Right, scatter plot of position along the anteroposterior axis (x axis) against the similarity with two choroid plexus *lipizones* (y axis), each point is a *lipizone* and is colored by its Allen Brain color. (**f**) Heatmap of the brain-wide correlation of sphingomyelins, highlighting three blocks (gray, white matter, and ventricular). The falling barplot at the bottom highlights the ventricular enrichment of each sphingomyelin. The red dashed line represents no enrichment (= 1.0). (**g**) Spatial plot of the four ventricular sphingomyelin markers, across the entire brain, for 5 representative sections. (**h**) Spatial zoom-in on the ventricles colored by the four ventricular sphingomyelin markers. (**i**) Barplot of the log2 fold changes of lipids in the ventricular systems compared to the rest of the brain. Sphingomyelins are highlighted in red (increased in ventricles) and orange (decreased). (**j**) Toxin staining validation of sphingomyelin in the choroid plexus. The toxins and the lipids they mark are indicated and color-coded. (**k**) Barplots of the Spearman’s correlations between lipid abundances and the dorsoventral axis in the lateral ventricle-lining *lipizone* (one bar per brain section), with (**l**) Spatial view of the lateral ventricle linings colored by individual lipids that were found to exhibit gradients along the dorsoventral axis.

We identified two distinct clusters in the choroid plexus (ChP), which we termed ChP1 and ChP2 (***Figure 5d, Supplementary Figure 10f***). While ChP1 was enriched in phospholipids and ether lipids, ChP2 contained relatively more HexCers (***Supplementary Figure 10g***). The differential lipid profiles suggested functional specialization: ChP2’s ceramide enrichment potentially indicated proinflammatory signaling capacity^57^, while ChP1’s phospholipid abundance suggested secretory functions^58^. Additionally, the two ChP *lipizones* aligned with ChP cellular composition, potentially corresponding to the ependymal-like cells (CHOR) and the non-ependymal components (e.g., mesenchymal and endothelial)^59^.

Fate mapping experiments demonstrated that roof-plate progenitors give rise to each choroid plexus from independent invaginations at four distinct roof-plate sites^60^. Moreover, bulk transcriptomics showed that adult choroid plexus retains ventricle-specific gene expression programs mirroring those embryonic origins^55^. We therefore wondered whether ChP1, enriched in lateral ventricles (***Figure 5e***), had a more anterior membrane lipid signature than ChP2, enriched in the fourth ventricle. We calculated the similarity between each of the two clusters and all brain *lipizones*, using Euclidean distance. At the same time, we tracked the average anteroposterior position of each *lipizone*, which revealed that *lipizones* located more posteriorly were significantly more similar to ChP2 (p < 0.0001 ; ***Figure 5e, Supplementary Figure 10h***). The only exception to this trend was a small cerebellar *lipizone* cluster, whose lipidomic profile was closer to ChP1. This suggests that the developmental positional identity may be retained in the choroid plexus lipidome.

Although sphingomyelins are typically classified as white-matter lipids, we found that they are unexpectedly enriched in the brain ventricles^13^. To explore this, we generated a whole-brain SM correlation matrix, which revealed three distinct clusters (***Figure 5f***): one broader and in the gray matter, with SM 36:2 and SM 36:1, metabolically linked with the neuronal marker Cer 36:1; one restricted to the white matter (42-long SMs); and another strictly ventricular (SM 34:1, SM 38:1, SM 40:1, SM 40:2). We propose SM 38:1, SM 34:1, SM 40:1, SM 40:2 as *bona fide* ventricular markers, as they visibly highlight the ventricular system in whole-brain views (***Figure 5g, h***; ***Supplementary Figure 10i***). We found that sphingomyelins, as a class, are vastly enriched in the ventricles compared to the rest of the brain (***Figure 5i***). Staining with toxins that specifically bind to SMs^61–63^ validated SMs as ventricular markers (***Figure 5j***). We identified six lipids (Hex2Cer 40:0, Hex2Cer 40:1, HexCer 42:2, HexCer 38:1;O2, PI 42:8), and, notably, the neuronal SM 36:1 showing an inverse trend, that displayed pronounced dorsoventral gradients along the walls of the lateral ventricles (|Spearman’s R| > 0.4; ***Figure 5k, l; Supplementary Figure 10j***). Since the walls of the lateral ventricles host the largest pool of neural stem cells in the adult brain^56,64^, which give rise to distinct neuron types depending on their dorsoventral position, the observed lipid gradients could potentially be related to differential neurogenic activity. Overall, our observations on SMs, the choroid plexus, and the ventricular walls reveal previously unappreciated lipid-based zonation within the ventricular system.

### Pregnancy reorganizes the brain lipidome

Mammals display sexually dimorphic behaviors that are associated with differences in neuroanatomical organization and circuitry between sexes^65^. Since bulk assays average local differences^66^, we turned to MALDI-MSI, characterizing the brain lipidome of 3 males and 3 females with approximately matched coronal sections (29 acquisitions in total among the 91 brain sections of the atlas; ***Supplementary Figure 12a***). The effect of sex on the lipidome was generally limited (Bayesian modeling detailed for the section on pregnancy; *see Methods*), with lipid levels having absolute logarithms of fold-changes (log2FCs) above 0.2 for a median of 1 lipid per supertype (***Supplementary Figure 12b***). Of the 222 supertypes, only 29 displayed a sex-related log2FC of at least 0.2 for more than 5/172 membrane lipids. A ventricular supertype however had 151 lipids all enriched in females, including moderate increases of phosphatidylcholines (PC 40:7, PC 42:1, PC 40:1; ***Supplementary Figure 12c***). We also assessed which lipids had recurrent sex-specific levels across supertypes, finding that Cer 40:2 was increased in females in 118/222 supertypes (***Supplementary Figure 12d***). On the other hand, the vast majority of lipids had highly-localized variation, with only 13/172 other lipids modulated in more than 10 supertypes by sex.

We next asked whether physiological events can instead dynamically modulate the lipidome. Pregnancy is a major physiological event, inducing metabolic adaptations across the organism^67^. Brain remodeling during pregnancy includes volumetric changes and circuit restructuring^68,69^ to support behavioral modifications and fetal metabolic demands^67,70–72^, which might involve lipid changes. We measured the brain lipidome of 3 first-pregnancy females (mid-gestation, embryonic day E13.5, ***Figure 6a***) by MALDI-MSI in addition to the 3 non-pregnant females, and we label-transferred 215/222 *lipizone* supertypes (***Figure 6b***). Embeddings analysis suggested that lipid variations are localized (***Figure 6c***) and overlapping with specific sets of *lipizones*, motivating us to characterize the response of individual *lipizones* to pregnancy. Using a Bayesian hierarchical model controlling for batch effects and inter-individual variation (***Supplementary Figure 12e-k***; *see Methods*), we quantified pregnancy-induced changes for each *lipizone* supertype (which we referred to as “shifts”, and the corresponding log2FCs, which were between −2 and +2), producing spatial shift maps that further confirmed that changes are spatially organized (***Figure 6d***). These changes could also be coarsely estimated (Pearson’s R = 0.54, ***Supplementary Figure 12l***) from a single brain section, highlighting the operational potential of the atlas as a reference and of the EUCLID label transfer function to detect localized lipid changes in low-data settings.

**Figure 6:**
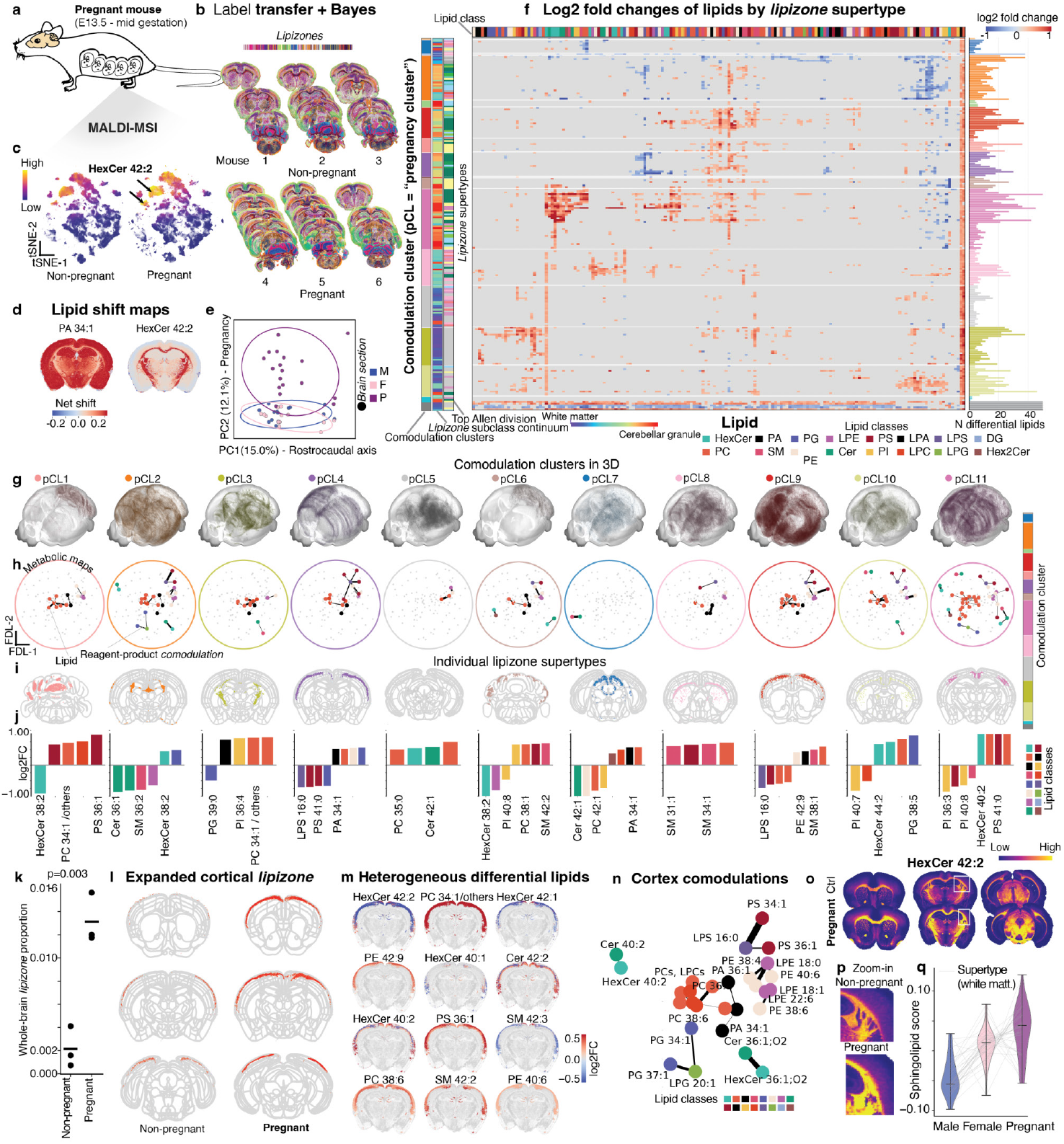
Lipidomic reorganization in the brain of pregnant mice. (**a**) Schematic illustration of spatial lipidomics analysis of pregnant mice (mid-gestation). (**b**) Spatial plots of the dataset used for the case-control hierarchical Bayesian analysis in pregnancy, colored by *lipizone* supertypes obtained with the label transfer algorithm. (**c**) tSNE, separated for pregnant and non-pregnant data, colored by the myelin marker HexCer 42:2. (**d**) Section colored by lipid shifts, i.e., the net difference upon pregnancy, for two recurrently-altered lipids in pregnancy. (**e**) Scatter plot of brain sections in PCA space (one point is one section), color-coded for males (M), females (F), and pregnant females (P). Ellipses capture the data ranges, and axes are labelled based on their correlations with the rostrocaudal axis and the pregnancy condition. (**f**) Heatmap (after optimal leaf ordering) showing the pregnancy log2FCs across all *lipizone* supertypes for all lipids, with leaf coloring indicating comodulation cluster (i.e., the clustering based on the pregnancy-driven changes), *lipizone* subclass, and Allen Brain division for the rows, lipid classes for the columns. The barplot on the right represents the count of differential lipids by supertype (outliers clipped). (**g**) 3D distributions of pixels belonging to each of the 11 pregnancy-driven comodulation clusters. (**h**) Metabolic maps as force-directed layouts (FDLs) highlighting comodulations between metabolically-connected lipid pairs, one plot per comodulation cluster. Dots are color-coded by lipid class; the layout structure is the same for all subplot; edge thickness is proportional to the comodulation score (see Methods). (**i**) Spatial plots of selected sections for individual supertypes, one per comodulation cluster, with (**j**) corresponding barplots showing their top increased and decreased lipids in response to pregnancy, colored by lipid class. The color coding is the same as in the remainder of the paper, e.g., *Figure 6f*. (**k**) Strip plot of *lipizone* proportion in non-pregnant and pregnant mice, for the cortical *lipizone* that was found to be largely increased in pregnant mice. (**l**) Spatial plot of the *lipizone* whose proportion was increased in pregnancy. (**m**) Spatial plots of some lipids with differential abundance upon pregnancy in the outer cortex, colored by their log2FC. (**n**) Metabolic map zoom-in for the comodulation cluster the increased-proportion *lipizone* belongs to. (**o**) Spatial plot of representative sections for control and pregnant brains, colored by HexCer 42:2. (**p**) Zoom-in on the white matter in a control and a pregnant brain section, colored by HexCer 42:2. (**q**) Violin plot of the sphingolipid score in males, non-pregnant females, and pregnant females, across core white matter *lipizone* supertypes. Individual *lipizone* supertypes are connected from male, to non-pregnant female, to pregnant-female with gray line segments.

We discovered an uncharted, organized, and vast lipidomic brain reorganization, suggesting that dynamic tissue needs can drive lipidomic heterogeneity. In fact, pregnancy induced more extensive changes in brain membrane lipids than sex differences between non-pregnant females and males of the same age (principal component analysis, ***Figure 6e***). The average change between a given supertype in pregnancy and control was 120% of the mean variation observed between sister supertypes (for the top 20 most altered supertypes), indicating substantial lipid remodeling during pregnancy, with an average of 16 lipids per supertype being modulated. Membrane score analysis (***Supplementary Figure 12m***), summing lipid log2FCs, revealed region-specific remodeling in pregnancy: negative scores (prevalent turnover) in ventricles, hindbrain, white matter, cerebellum, outer cortex and hypothalamus, and positive scores (prevalent biosynthesis) at gray-white matter boundaries and in deep cortical layers.

The lipidomic reorganization induced by pregnancy involved 139/215 supertypes (as having > 5% differential lipids among their respective expressed lipids; ***Figure 6f***). We found that some lipids were recurrently altered, such as HexCer 42:2 and PA 34:1, but overall differential lipids were highly heterogeneous across supertypes (14 affected supertypes in median across lipids, log2FC > 0.2 or < −0.2 for >98% posterior samples), suggesting adaptations to pregnancy dictated by local needs. To make sense of this complexity, we clustered supertypes based on their log2FCs across lipids, highlighting 11 “comodulation” clusters (*see Methods*) that modulated the lipidome differently from one another in response to pregnancy (***Figure 6f, g***). These comodulation clusters were aligned with the *lipizones* hierarchy and the Allen Brain divisions. Lipid modulation also seemed to be broadly organized by lipid class, even if less clearly; thus, to better understand the metabolic axes of modulation, we calculated a comodulation score for metabolically-connected lipid pairs (see Methods; ***Figure 6h***). This score provides a first indication of the reactions likely modulated in pregnancy. Modulated metabolic pairs varied widely among clusters, from one or few to entire subnetworks (eg, the network involving PS 42:6, PE 42:6, LPE 22:1, and more in the “pink” comodulation cluster), encompassing multiple types of enzymatic activities (eg, enzymes *Pla2g2a* and *Ptdss2*). These observations further supported the view that pregnancy entails localized modulations. In every comodulation cluster, several supertypes were altered, with specific profiles that do not trivially reflect lipid classes only (***Figure 6i, j***). Driven by the specificity of these changes, we characterized the response to pregnancy for individual supertypes.

First, we identified a supertype localized in the cortical layer 1 that expanded dramatically during pregnancy, from rare to an 11-fold higher representation (p = 0.002, tested across individuals), with significant modulation at absolute log2FC 0.2 of 16 lipids (***Figure 6k, l***). Overall, the outer cortex as a region exhibited vast lipid remodeling − 169/172 lipids were significantly changed, of which 40 increased and 23 decreased with sizeable effect size (absolute log2FC > 0.2; Wilcoxon test), including changes in HexCer 40:1 (increased) and PC-O 32:0 (decreased) (***Figure 6m***). The cortex is in fact known to be vastly reshaped by pregnancy in humans^73^; in mice, cortical changes in pregnancy were related to maternal behavioral functions, with the females exhibiting more maternal behaviors showing temporary cortical hypertrophy^74^. The negative membrane score indicative of prevalent turnover and the heterogeneity of comodulations (involving PCs, PGs, PEs, PAs, PSs, and sphingolipids; ***Figure 6n***) suggest that functional changes in the outer cortex are paralleled by complex lipidomic rearrangement.

Finally, we found that the myelin marker HexCer 42:2 was vastly increased by pregnancy across core white matter supertypes, in all subjects (average log2FC across supertypes = 0.38; ***Figure 6o, p, Supplementary Figure 12n***). Sphingolipids were also collectively modulated, with a prevalent upward trend (***Supplementary Figure 12o***). This observation suggests widespread adaptive myelination by pre-existing oligodendrocytes as oligodendrogenesis in pregnancy is supposed to be more restricted in space compared to the large-scale HexCer 42:2 change^75^.

Sphingolipids also had a prevalent upward trend from males to females in the white matter; therefore, we computed a sphingolipid signature, which increased across white matter supertypes from males, to non-pregnant females, to pregnant females (***Figure 6q***). Collectively, these observations identify membrane lipid composition as an axis that can be modulated in response to physiological events beyond inter-individual variation, producing highly structured, regionalized responses.

## DISCUSSION

Using high-resolution MALDI-MSI and computational methods, we mapped the lipidomic architecture of the adult mouse brain. We identified over 500 biochemically distinct territories that align with anatomical regions and correlate with cell types and their connections. These biologically meaningful territories, which we termed *lipizones*, represent the fundamental units of lipid composition, and they expose previously unrecognized metabolic organization.

The relationship between brain membrane lipid composition and cell type distribution emerged clearly in our study. We identified 304 *lipizones* that overlap with transcriptionally-defined cell types, revealing a connection between the cellular and the lipid architectures. Importantly, *lipizones* were able to capture cell bodies and their distant axon terminals, suggesting their potential use to support the investigation of a dimension of the brain architecture not measured by spatial transcriptomic methods.

Classical neuroanatomical boundaries are defined by cytoarchitecture and connectivity, but their relationship to metabolic heterogeneity was unknown. We discovered sub-anatomical zonation in the cerebellar granule cells layer and callosal axon bundles, indicating that cytoarchitecture and myelin content create biochemical substructures within classical boundaries. We showed that connected brain regions, even those physically distant, share distinct lipid signatures. This observation suggests that lipidomic profiling could contribute to building a scalable label-free proxy for connectivity mapping, and positions lipids as an important new layer to interrogate brain function beyond gene expression.

Myelin has traditionally been characterized as relatively homogeneous across the brain, and transcriptomic studies have identified limited oligodendrocyte diversity. In contrast, our lipid analysis reveals striking regional diversity in white matter composition, predominantly driven by oligodendrocyte-specific lipids. We attribute part of this diversity to the chronotopic unfolding of myelin development, an explanation compatible with the reported stability of myelin.

Finally, during development, maternal metabolism changes in response to the fetus and to adapt behaviorally to the postpartum. We found large-scale, coordinated lipid adaptations in the brains of pregnant females, with changes in the outer cortex and white matter production. These results demonstrate the utility of *lipizones* for future studies on both normal physiology, such as aging and dietary effects, and disease conditions.

Our observations suggest that females have higher myelin production capacity than males, further boosted by pregnancy. Major depressive disorder (MDD) modulates the lipidome^76^, with depressed female adolescents showing increased myelin in white matter tracts^77^; pregnancy is positively associated with depression^78^. Furthermore, myelin sphingolipids may contribute to women’s higher susceptibility to Alzheimer’s disease and demyelination^79,80^. Disease-related stressors might over-activate this already enhanced myelination system in females, resulting in larger-scale white matter changes.

The lipidomic heterogeneity we uncovered provides a compelling foundation for future functional studies. For instance, the enrichment of sphingomyelins in ventricles and the unexpected diversity in white matter composition suggest specialized functional roles yet to be characterized. Similarly, in pregnancy, distinguishing between adaptations supporting maternal behavior and those meeting fetal metabolic demands could reveal specific regulation mechanisms of the maternal-fetal neurometabolic interactions. These examples highlight the rich potential for functional discoveries within the lipidomic architecture we have mapped.

## METHODS

### Animal work and cryosectioning

Healthy adult mice (approximately 8 weeks old) were anesthetized using isoflurane and euthanized by cervical dislocation. Brains were dissected, immediately embedded in 2% carboxymethylcellulose (CMC) in deionized water, and frozen on dry ice plus isopentane. Samples were stored at −80°C until cryosectioning, where coronal sections were cryo-sectioned with a cryotome into 10 µm sections. Sections were collected at 200 µm intervals throughout the entire brain. For sex and pregnancy comparisons, six representative regions were sampled at approximately the following stereotaxic coordinates (in mm, relative to interaural and bregma): 1) 5.14; 1.34; 2) 3.22; −0.58; 3) 2.34; −1.46; 4) 1; −2.80; 5) −0.92; −4.72; 6) −2.84; −6.64. All animal care and treatment procedures were performed in accordance with the Swiss guidelines and were approved by the Canton of Vaud SCAV (authorization VD 3730.c).

### MALDI-MSI experimental acquisition

Sections were dried at room temperature and coated with 2,5-Dihydroxybenzoic acid (DHB) (30 mg/μL in 50:50 acetonitrile:water and 0.1% trifluoroacetic acid) using the automatic SMALDIPrep (TransMIT GmbH) at 350 rpm and flow rate 5 μL/min for 30 min. Acquisitions were performed with an AP-SMALDI5 AF system coupled to a Q Exactive orbital trapping mass spectrometer. Calibration maintained mass error within ±2 ppm. For each pixel, the spectrum was accumulated from 50 laser shots at 100 Hz. MS parameters in the Tune software (Thermo Fisher Scientific) were set to the spray voltage of 4 kV, S-Lens 100 eV, capillary temperature to 250°C. The step size of the sample stage was set to 25 μm. Positive ion mode measurements were performed in full scan mode (m/z 400-1600) with resolving power R = 240000 at m/z = 200, using internal calibration via lock mass.

### LC–MS/MS Analysis

Four male and four female C57BL/6J mice (Charles River) were euthanized with CO2 and decapitated. Brains were rapidly extracted and snap-frozen in liquid nitrogen, then ground in a pre-cooled mortar while continuously adding liquid nitrogen until reaching a homogenous powder (∼5 min). The powder was aliquoted and weighed to determine wet weight. Brains were analyzed by high-throughput targeted HILIC MS/MS lipidomics. Complex lipids were extracted with isopropanol pre-spiked with internal standards containing 75 isotopically labeled lipid species. Extracts were analyzed by hydrophilic interaction chromatography coupled to electrospray ionization tandem mass spectrometry (HILIC ESI-MS/MS) in positive and negative ionization modes using a TSQ Altis LC-MS/MS system. Using a dual-column setup, multiple lipid classes (glycerolipids, glycerophospholipids, cholesterol esters, sphingolipids, and free fatty acids) were quantified. HILIC separation allowed endogenous lipids to co-elute with corresponding internal standards for matrix effect correction. Data acquisition used timed selected reaction monitoring (tSRM) mode with lipid class-dependent parameters. Raw data were processed using Trace Finder Software (v4.1) with abundance reported as estimated concentration.

### Registration to Allen Brain Common Coordinate Framework

We registered to the Allen Brain Atlas (ABA) common coordinate framework (CCFv3^19^). First, manual registration with Aligning Big Brains and Atlases (ABBA^81^) software employed affine transforms and nonlinear warping with anatomical borders guiding key feature registration. For improved accuracy, we further processed sections with STAlign^82^, using the Allen density images^83^ as reference and peak m/z = 845.528 as target (which is the most correlated with density). The resulting diffeomorphism was interpolated to reposition lipidomic pixels.

### Raw data processing with uMAIA

We used uMAIA^18^ (github.com/lamanno-epfl/uMAIA/) with default settings to process all raw data collectively (peak calling, peak matching, feature normalization), including two complete individual male brains (73 sections), male and female control sections from 3 individuals (29 sections), and brains of 3 pregnant mice (18 sections). We identified 26,874 features initially. Quality control restricted the feature space to [M+0] isotopes, biological compounds (greater intensity within tissue than matrix), and features present in ≥4 consecutive sections. Before normalization, we reduced the feature space by removing peaks not annotated in any database, peaks with Moran’s I <0.4, and redundant adducts. For redundant adducts mapping to the same lipid, we kept the peak with the least dropout and the highest total signal after confirming a high correlation between different adducts on non-dropout sections. This procedure yielded 1,400 m/z peaks for normalization with uMAIA.

### Data preparation and feature selection

We extracted uMAIA-normalized m/z peak values for all pixels and acquisitions, transformed them back to linear scale, and stored them in a dataframe with spatial coordinates and Allen brain metadata (anatomical regions, colors, contours). We refined tissue masks using CCF borders, removing pixels mapping outside the brain volume. To completely remove features (m/z peaks) with residual batch effects, we performed the following feature selection procedure. We computed three scores: (1) difference between intra- and inter-acquisition variance, (2) spatial autocorrelation computed as Moran’s I (squidpy^84^), and (3) number of sections the lipid measurement dropped out. First, we removed features with dropout > 4 sections (retaining 247/1400 features), then we clustered lipids in the space of these scores (standardized) with kMeans (k = 10), and removed the features belonging to 5/10 clusters, corresponding to visibly corrupted distributions across acquisitions, retaining 105/247 features.

### Dimensionality reduction and batch integration

All differential testing and downstream analyses of the manuscript were performed directly on the uMAIA output. The dimensionality reduction and harmonization described below was used solely for clustering where strong coembedding is desired.

Dimensionality reduction was performed fitting non-negative matrix factorization (NMF) on a brain #1, providing interpretable “global” embeddings.

Non-negative matrix factorization (NMF) seeks two low-rank non-negative matrices, 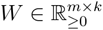 and 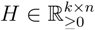, whose product approximates the original non-negative data matrix: 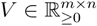

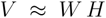

Here, *m* is the number of pixels, *n* the number of lipids, and *k* the number of NMF factors. The number of NMF factors was selected as the number of lipid Leiden clusters maximizing the score:

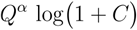

where Q is the modularity, which quantifies the strength of division of a network into clusters, C the number of clusters, and α a tunable parameter that controls the relative influence of modularity versus having a large number of clusters (default = 0.7). This yielded 16 lipid Leiden clusters, i.e., 16 NMF factors. The NMF component matrix (left-hand side, W matrix) was seeded^22^ with the lipid closest to the centroid for each cluster of lipids and the coefficient matrix with the zero-clipped cosine similarity between each lipid and the central one. We verified embedding robustness to random feature downsampling. Once the W and H matrix were learnt from brain #1, the H matrix was kept fixed and the W matrices were determined for the other brains; Harmony^23^ was applied on the W matrices. We also produced a low-rank, batch effect-corrected dataset by multiplying the component with the coefficient matrix. This was used for the local embedding recomputation during the clustering procedure described below. The coefficient matrix was z-standardized.

### Top-down binary splitting algorithm and label transfer

We implemented a top-down hierarchical binary splitter. The algorithm splits the set of input pixels in two recursively. To perform each of the split balancing local and global variations, the globally computed H matrix (NMF coefficients for each pixel) is concatenated with the H matrix of the pixels of the parent split and of the current split, i.e., the NMF is iteratively recomputed at each split. The optimal binary split is found using the backSPIN^85^ approach, which is sped up by using aggregated pixel clusters instead of individual pixels (k=60 clusters using kMeans).

A stop condition is implemented so that the splitting is halted if one of the following conditions is not satisfied: (i) differentially expressed features (inspired by Yao et al^1^), (ii) minimal size of clusters and (iii) coherence between consecutive sections. For differential expression, Mann-Whitney U-test with BH FDR-correction required ≥2/105 differential peaks at 0.2 absolute fold change and 0.05 adjusted p-value. The minimal size of a cluster was set to 200 pixels. For coherence between consecutive sections, we impose that each cluster must span at least 2 consecutive or 4 total acquisitions.

To make the split robust (i.e. not instance-dependent) and to allow for accurate label transfer of new datasets, at each split we trained an XGBoost binary classifier to perform the split from the embeddings. XGBoost classifiers (class rebalancing, 1000 estimators, max depth 8, learning rate 0.02, 0.8 subsampling and colsample, and gamma 0.5) used early stopping on validation set (i.e., a set of held-out sections spanning the anteroposterior axis) accuracy. The XGBoost classifications on the training, validation, and test sets were used as cluster labels. By this procedure, our clustering method inherently builds a transferrable annotation tree: a new pixel can be annotated by applying the local NMF to obtain its embeddings, followed by XGBoost classification, moving iteratively from the root to a terminal leaf (*lipizone*) of the binary tree.

### *Lipizone* assessment, naming, and visualization

We compared our clustering approach with Leiden clustering (resolution=14.0) inspecting the confusion matrix. Results were qualitatively confirmed by visualizing with matching colors the clusters. Further assessment of stability was performed by clustering independently brain #2, retrieving clusters in good agreement with those label-transferred from the clustering of brain #1.

*Lipizones* were named based on the Allen Brain anatomical acronyms where they were found. Specifically, we calculated a reciprocal enrichment score from the confusion matrix of *lipizones* and Allen brain acronyms by element-wise multiplication of the enrichment of *lipizones* in acronyms and the enrichment of acronyms in *lipizones. Lipizone* colors were chosen to allow the visualization of multiple lipizones in the same view. To maximize contrast, we colored the *lipizones* concatenating 8 colormaps, one for each of the 8 *lipizone* classes, and assigned the colors based of the lipizone “leaves” of the hierarchy. Contiguous leaf *lipizones* were distanced along the colormap proportionally to their cosine similarity.

### Feature restoration

We used XGBoost regression to impute features discarded during selection (i.e. the ones where measurement dropped out in a non negligible numer of sections). Features with Moran’s I ≥0.4 in ≥4 sections were candidates for restoration. For each feature, we selected the 3 best sections by Moran’s I for training and the fourth for testing. XGBoost models were trained to predict discarded features from the 16 Harmony-corrected NMF embeddings. Models yielding Pearson’s R ≥0.4 on held-out test sections were accepted and applied to all sections, resulting in 692 m/z peaks restored.

### Lipid annotation

For an accurate annotation of m/z peaks, we integrated multiple reference sources: three bulk lipidomic experiments (ESI LC-MS with two different chromatographic column, HILIC and Reverse Phase, and a LC-MS/MS), METASPACE annotations (using the Human Metabolome Database - HMDB), the LIPIDMAPS database, and two published brain MS datasets (Fitzner et al.^11^ LC-MS and Vandenbosch et al. MALDI-MSI^14^). We matched raw peak m/z values to reference annotations within 5 ppm, harmonizing nomenclature with goslin^86^ when needed. For MALDI-MSI, we considered [M+H]+, [M+Na]+, [M+K]+, and [M+NH4]+ adducts; for in-house LC-MS/MS, we used [M+H]+ and [M+NH4]+ in positive mode and [M+AcO]−, [M-2H]−, and [M-H]− in negative mode.

We assigned confidence-weighted scores to each annotation: LC-MS/MS (8), ESI LC-MS (2), published studies (1), METASPACE (1 if FDR<0.05, otherwise 0.5), and LIPIDMAPS (0, annotation transferred but unconfirmed in brain). Peak names were further prioritized using an independent quantitative LC-MS/MS dataset for sex-specific brain lipidomics (***Supplementary Table 1***). In cases of naming ambiguity, we prioritized lipid names covering >80% molar fraction among candidates; otherwise, we reported multiple names as the species designation. This approach allowed us to disentangle several PE and PC isomers while flagging some peaks as potentially representing multiple isomers. We provide detailed annotation tables (***Supplementary Table 2a, b***) motivating every lipid name where an overlap of an abundant and a less abundant species is possible.

Peaks with annotation scores >4 (172 nonredundant lipids) were considered unambiguous and used for the analysis in this work. The final dataset includes multiple lipid classes (LPC, LPE, LPA, LPS, LPG, LPI, PC, PE, PI, PS, PA, PG, SM, Cer, HexCer, Hex2Cer, TG, DG) with variable chain lengths and unsaturation degrees. Since MALDI-MSI cannot distinguish isomers, we report the sum composition of carbon atoms and unsaturations, meaning an “individual” lipid may comprise multiple isomers. We created a lipid property table using regular expressions to extract class designations, carbon numbers, and unsaturation counts.

### Data analysis and visualization methods

For visualization, we embedded our dataset in t-SNE space (perplexity 30) using z-standardized, harmonized NMF embeddings.

3D interpolation used radially exponentially-decaying weighted neighbors with Allen brain regions as anatomical guides. To render, explore, and film 3D maps, we used Napari^87^ and napari_animation (***Supplementary file 1***).

We assessed the residual within-cluster variability of each *lipizone* on brain #1 as the log determinant of the covariance matrix of that *lipizone*, after z-scaling lipids across the entire brain. We processed cell type-specific bulk lipidomics data from Fitzner et al.^11^ using min-max normalization. For the oligodendrocyte score used in the white matter analysis, cell type deconvolution used linear regression by non-negative least squares, with coefficients normalized to sum to 1 across cell types per *lipizone*. Differential lipid testing between groups used Mann-Whitney U-test with Benjamini-Hochberg correction (α=0.05). A permutation test was used to evaluate lipid class enrichment.

For the similarity range score comparing the two ChP major *lipizones* (ChP1-ChP2) against all other brain *lipizones*, for each *lipizone* centroid, we computed the Euclidean distances to both ChP *lipizone* centroids in lipidomic space restricting to differential lipids. We then expressed this as a relative distance ratio, where values below 0.5 indicated greater similarity to the first ChP *lipizone* and values above 0.5 to the second.

### lipiMap model and Lipid Ontology integration

lipiMap is a Variational Autoencoder (VAE) adapted from expiMap^44^. The encoder employs a feedforward neural network (four hidden layers: 256, 256, 256, 128 neurons) with layer normalization, ReLU activation, and dropout. The decoder is a single linear layer with weights masked by lipid membership in specific lipid programs (LPs), in order to yield a biochemically interpretable latent space. Input data was stretched to the range [0, 1000] and approximated to the next integer. Training used brain #1 (1.7M pixels) with 10% held out for validation and male/female control brains as the test set. We implemented the network using PyTorch (version 2.2.1), and we trained the model on a single NVIDIA GeForce RTX 4090 GPU. Optimal hyperparameters were determined by random search over the hyperparameter space, and the best-performing configuration was selected by choosing the model with the lowest ELBO loss on the validation set. The Adam optimizer^88^ was used to optimize the ELBO loss function during training.

To constrain the latent space with biochemical knowledge, we downloaded Lipid Ontology^45^ (LION v2020-08-07) from BioPortal, using four categories of LPs: lipid classes, cellular component, functional categories, and physicochemical properties. For example, some LPs include:

- The lipid class “PC” (phosphatidylcholines)
- The cellular component “mitochondrion”
- The functional category “lipid-mediated signaling”
- The physicochemical property “high transition temperature”

165/172 lipids appeared in more than one LP and were used for downstream analysis. To assess whether each lipid program was sufficiently represented by the lipids detected in this study, we performed bootstrap analysis with 10,000 random samples from 1,176 brain lipids found in LION, calculating a representation score for each program. Specifically, we defined the representation score as the ratio of annotated lipids to total lipids associated with each LION term. Programs with representation scores below bootstrapped median or <3 lipid members were excluded. The matrix of lipid to program membership was used as the mask for decoder weights, enforcing interpretability by implementing hard lipid membership to LPs. This means that membership cannot be learnt during training, but it is fully pre-defined, using LION. The decoder weights matrix was initialized using PCA loadings and masked with the LP matrix. The tendency of VAEs to generate outputs that regress towards the mean of all inputs was mitigated through a weighted MSE reconstruction loss for training:

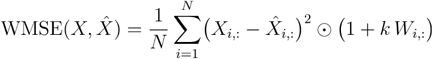

where W_ij_ = 1 if X_ij_ is an extreme value, W_ij_ = 0 otherwise, and k is a tunable parameter. Briefly, for each lipid species, extreme values were defined as those below its lower q-th quantile or above its upper (1-q)-th quantile, with quantiles precomputed across the entire training dataset. We enforced a direct relationship between the sign of latent coordinates and their contribution during linear reconstruction by the decoder, specifically, negative latent coordinates should correspond to lower associated lipid levels, while positive coordinates should lead to higher. To obtain this behavior, decoder weights were clipped within the range [0, 1], so that a deactivated LP (i.e., negative latent coordinate) cannot combine with a negative weight to paradoxically produce a positive contribution.

### Multi-omics integration

We used a whole-brain MERFISH dataset (from Yao et al.^1^ - and the 8460-genes imputed dataset provided by the same paper) to assess whether some *lipizones* overlapped spatially with cell type territories and to compare lipid and gene expression distributions. This dataset was warped into the same coordinate framework as the Lipid Brain Atlas, namely, the Allen Brain CCFv3. For each pixel of the Lipid Brain Atlas, we evaluated whether any cell was available in the MERFISH dataset within a neighborhood of radius of 50 μm in 3D, after filtering out immune and vascular cells. If at least one cell was available, a cell type territory match was putatively assigned to the pixel by majority voting across the cells of the neighborhood. For gene expression, we averaged across the cells in the neighborhood. To avoid spurious matches across the main anatomical borders due to imperfect registration, we allowed neighborhood to be established only within the main anatomical regions. Next, to filter well matched sections, we evaluated the overlay between gene *Mog* and lipid HexCer42:2, as both mark myelin. The resulting 53 sections were uniformly distributed along the anteroposterior axis. Analogously, we considered a Slide-seq2 whole-brain dataset (from Langlieb et al ^3^) to validate our results. To directly compare with this dataset, we used STAlign^82^ to warp the Slide-seq2 sections onto the Lipid Brain Atlas matched sections (assessed by physical proximity in CCF space), using the Slide-seq2 Nissl images. We retained ∼120k Slide-seq2 beads marked as not containing a doublet of cells, and generated an imputed full transcriptome (21,877 genes) using the snRNA-seq data from the same study. The lipidome was evaluated on each Slide-seq2 bead as the profile of its closest MALDI pixel. Next, for each cell type, we averaged the lipidome of the beads belonging to that cell type.

### Prediction of lipid levels from gene expression

To predict lipids from genes, we trained linear models and XGBoost regressors using mean squared error as the loss, using spatial pixels with MERFISH data and cell type averages with Slide-seq2 data. We trained a canonical correlation analysis (CCA) model on 538 MERFISH-imputed metabolism-related genes and our 172 lipids. Redundancy was calculated as the mean squared Pearson’s R between each canonical axis derived from the CCA and each lipid. This metric represents the fraction of lipidomic variance captured by each gene-derived canonical variate, and summing across components yields the cumulative redundancy, indicating the total variance in the lipidome explained by the leading gene–lipid association modes.

We also trained XGBoost models to predict a leave-1-out lipid from all the other lipids, with the same parameters of gene to lipid models, in order to estimate the irreducible fraction of variance *F* for each lipid, where:

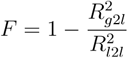

with R_g2l_ is the Pearson’s R between ground truth and reconstruction for the genes to lipid prediction and R_l2l_ refers to the same score for the lipids to leave-1-out lipid prediction of the same target lipid.

### Feature importance analysis on full transcriptome

Cell type-specific gene enrichment was calculated as previously described in Zeisel et al.^89^ as:

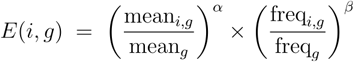

across the 8460-dimensional MERFISH-imputed transciptome dataset. Mean and freq represent expression level and detection frequency for gene *g* in cell type *i* versus all cells, defaulting to α=β=1. Genes were assigned to cell types based on maximum enrichment scores above a defined threshold, excluding vascular and immune populations.

We trained an XGBoost regressor for each of 172 lipids on the z-scored 8,460-dimensional transcriptome (predictor). To characterize the genes that contribute the most to predicting lipids, we extracted the SHAP feature importances^43^ for each gene, for each lipid prediction, on every pixel. SHAP values were then averaged within 251 Allen Brain Atlas regions (requiring minimum pixel coverage), converted to absolute values to capture feature importance magnitude, and normalized for cross-lipid comparability. To assess whether genes with the highest feature importances for lipids prediction were enriched for known cell type markers, we performed a permutation test (B=2,000) on the top 20 SHAP-ranked genes per lipid-region pair. For each pair, we compared the mean marker rank of the top 20 genes against a null distribution generated by randomly sampling 20 genes without replacement. Raw p-values were corrected for multiple testing using Benjamini-Hochberg FDR (α=0.05).

### Colocalization of *lipizones* and cell type territories

*Lipizone* overlap with cell type territories was defined as *lipizone*-cell type colocalization in terms of reciprocal enrichment (the enrichment of *lipizones* in cell types multiplying the enrichment of cell types in *lipizones*).

Specifically, let:

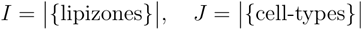

and for lipizone *i* and cell-type *j* let:

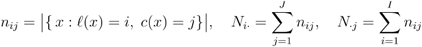

Then the two directional enrichments are:

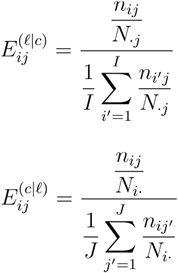

and the reciprocal enrichment score is the element-wise multiplication:

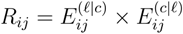

A threshold of reciprocal enrichment (= 200) and a threshold on the number of pixels where a given cell type and a given *lipizone* were overlapping (= 100) were chosen empirically to decide that a *lipizone* matches a cell type territory.

### Connectomic data and statistical analysis

To test whether highly connected region pairs share on average at least one *lipizone* above chance levels, we integrated three matrices: the ipsilateral connectomic strength matrix (C), a physical distance matrix (D), calculated from the Allen Brain Atlas regional centroids, and a region by region *lipizone* similarity matrix (S). C was downloaded from Oh et al.^4^; S was obtained as, where X is the binarized colocalization matrix between *lipizones* and Allen anatomical acronyms using reciprocal enrichment as the score. Highly connected pairs were defined as those exceeding the 95th percentile of C, and the observed test statistic (T_obs) was computed as the difference in the proportion of pairs sharing at least one *lipizone* between highly connected vs other pairs. To account for the confounding effect of spatial proximity, we estimated the conditional probability of high connectivity as a function of physical distance by binning D and computing the fraction of pairs meeting the connectivity threshold within each bin. Connectivity labels were then resampled for each pair from a Bernoulli distribution parameterized by this estimated probability, to preserve the marginal distribution of inter-regional distances. Repeating this procedure for 1,000 iterations generated an empirical null distribution for T, against which T_obs was compared using a one-tailed test; the p-value was calculated as the fraction of simulated T values that equaled or exceeded T_obs. The same qualitative results held when using the contralateral, instead of the ipsilateral, connectivity matrix, and when varying binning and connectivity thresholds.

### Integration of cell type and connectomic data to analyze cell terminals

We devised a strategy to assess whether a *lipizone* might contain both the cell bodies and the terminals of a neuronal population. We focused on the *lipizones* that were detected as substantially overlapping with one or more cell type territories based on the MERFISH dataset. We extracted the pixels belonging to a given *lipizone*, and clustered them in the space of their 3D spatial coordinates using kMeans clustering with k = 2. For each *lipizone*, we then asked whether one of these two *lipizone* subclusters was largely responsible for the overlap with the corresponding cell type territory in the MERFISH data. Since MERFISH is a nucleus-based technique, a *lipizone* spatial subcluster overlapping > 80% with the corresponding MERFISH territory suggests a cell body subcluster. To further support the observation that a *lipizone* might contain two spatial subclusters representing the cell bodies and the terminals of a cell type, we looked for axonal streams connecting these two subclusters. To do so, we used the whole-brain mesoscale Allen connectome^4^. This connectome, downloaded via API using the MouseConnectivityCache, traces axons from 2000+ viral tracer injection sites through the entire volume of the mouse brain at submicrometric resolution. Every tracer injection site is available as a different channel, and it represents a connectivity stream. We binarized the connectome intensities (signal above 95-percentile, independently for each channel) to capture whether a stream passes through a pixel or not, and we evaluated the connectome on the positions where we measured a lipidomic pixel. *Lipizone* spatial subclusters were putatively assigned as containing cell body and terminal territories when 1+ connectomic streams passed through both spatial subclusters, with the two subclusters at opposite extremes of the stream, as assessed using the first eigenvector of the spatial coordinates of all points spanned by the stream.

### Analysis of white matter clusters in neuronal and oligodendrocyte feature spaces

We ran three computational experiments to determine whether oligodendrocyte or neuronal lipids (from Fitzner et al.^11^) drive white matter heterogeneity. First, we retrained a 65-class XGBoost classifier using as the only sets of features the oligodendrocyte-restricted and neuron-restricted lipids. We then used SHAP to extract class-wise feature importances for each lipid, and assessing the relative enrichment of the neuronal-restricted and oligodendrocyte-restricted lipids in the top predictive features set by gene set enrichment analysis (GSEA)^90^.

Second, we subset the pixels belonging to the 65 white matter *lipizones*, and we reclustered, independently on the neuron-specific and oligodendrocyte-specific lipid sets, these pixels using N=10 principal components to avoid feature number imbalance, and proceeding with Leiden clustering. We next compared the 65 *lipizones* with the clusters deriving from these restricted feature spaces computing the confusion matrix, NMI and ARI scores.

Third, we retrained a 65-class XGBoost classifier as above by using as features the “lipid programs” inferred from our lipiMap model, i.e., its latent space, and extracting the feature importances using SHAP.

### Case-control analysis method development

We formulated a hierarchical Bayesian model to analyze changes during pregnancy or dictated by sex, while accounting for *lipizone* supertypes and potential batch effects. The model incorporates four nested biological variation levels with the following likelihood:

For lipid abundance *x*_*gst*_ at pixel *g* of supertype *t* in section *s*:

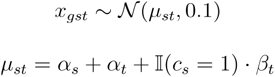

where *c*_*s*_ indicates the condition for section *s*. The parameters *α*_*s*_ and *α*_*t*_ represent section and supertype (“baseline”) effects, respectively, while *β*_*t*_ captures the condition effect for supertype *t* (“shift”). The indicator function I(*c*_*s*_ = 1) is 1 if section *s* is in the pregnant condition, and 0 if it is in the control, non-pregnant condition. The model structure is as follows:

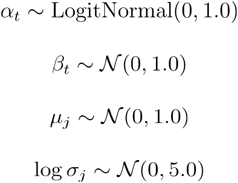

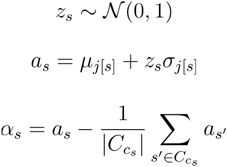

where j[*s*] maps section *s* to its parent sample j. We impose a sum-to-zero constraint: where C*c*_*s*_ denotes sections in condition *c*_*s*_. Any global, systematic offset of all pregnant sections versus all non-pregnant ones cannot be distinguished from a batch artifact a priori. By enforcing that the mean section effect is zero in both conditions, we guarantee that the model attributes any consistent difference to the biological “shift” term. Meanwhile, allowing each sample to have its own section-level standard deviation captures the fact that batch effects manifest at the level of individual slides rather than entire samples.

For inference, we employ a mean-field variational family:

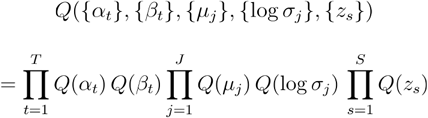

The variational distribution is parametrized as follows:

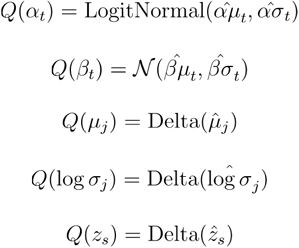

To prevent spatial autocorrelation to overinflate posterior confidence, data was prepared by ensuring that only one pixel within each 3x3 squared neighborhood would be in the training (and in the test) set, selecting pixels that respected the criterion randomly until pixel exhaustion. The model was implemented in NumPyro^91^ and fit using stochastic variational inference, estimated through 2,500 Adam optimizer iterations (learning rate=0.01) minimizing the ELBO loss.

Performance evaluation included loss convergence; plateau of parameter traces; Pearson’s correlation between measured and reconstructed supertype centroids by section on training and test pixels; QQ-plots of observed versus posterior predictive distributions; and parameter-stability tests comparing estimated shifts to simple centroid differences. For inter-individual variation detection, we compared the inferred sample means with the empirical sample averages. We evaluated 15 hyperparameter configurations (varying downsampling ratio, random seed, learning rate, number of epochs, prior scales and guide initialization) on ten lipids spanning a range of batch-effect severities, exporting visualizations and inspecting each posterior distribution one parameter at a time. Learning curves over 2,500 epochs were assessed to confirm that posterior means and uncertainties reached stable plateaus, with only modest uncertainty increases under downsampling. Pixel-level XGBoost models Pearson’s Rs between ground truth and fitted, trained on female samples (pregnant and non-pregnant), correlated strongly (Pearson’s R = 0.73) with our hierarchical model’s performance in terms of Pearson’s R, validating its prediction abilities. log2FC was defined as the log2 of the ratio of a supertype baseline plus its shift and the baseline. Significantly shifted lipid-supertype pairs were detected as having >=98% posterior samples with log2FC > 0.2 or >=98% posterior samples with log2FC < −0.2.

### Reaction matrix

We constructed a reagent-product-enzyme table using LINEX2^30^, followed by manual literature-based curation (***Supplementary Table 4***). We defined sixteen biochemical criteria as logical conditions to capture biologically plausible lipid transformations, including: LPX↔PX conversions (maintaining the “X” moiety), PC→PA transitions, PE ↔PC, PE↔PS, LPC→longer-chain LPC, and sphingolipid interconversions (SM↔Cer↔HexCer). These conditions were implemented via string operations and boolean masking, filtering only lipid-lipid pairs meeting at least one criterion. This computational approach yielded putative reactions that matched 100% with manually curated LINEX2 extractions. Each reaction pair was assigned its putative enzyme using a curated lookup dictionary mapping reagent-product combinations to enzymes (e.g., Lpcat1/2 for LPC→PC, Pla2g2a for PC→LPC, Smpd1-4 for SM→Cer).

### Lipid-lipid comodulation analysis in pregnancy

For each of the 172 annotated lipid species, we extracted the Bayesian model parameters for the shift and supertype baseline, and generated 1000 posterior samples using normal distributions with these learned location and scale parameters. Modulation was defined as shifts exceeding 20% of baseline expression. Lipids were classified as increased, decreased, or none when >98% of samples had shift > 0.2×baseline, or shift < −0.2×baseline, respectively. Expression was evaluated as baseline > 0.05 for >98% of posterior samples. For each supertype, we calculated the proportion of expressed lipids showing significant regulation (increase or decrease) and computed 95% confidence intervals using bootstrap resampling (10,000 iterations) to assess the reliability of regulation estimates across supertypes. Lipid shits were transformed into log2 fold changes compared to non-pregnant baseline, and hierarchically clustered using cosine distance with a custom metric that handles sparse data by excluding zero-zero pairs and returning maximum distance for zero-norm vectors. Clustering used weighted linkage with optimal leaf ordering, and supertypes were partitioned into 16 clusters using the fcluster algorithm (“comodulation clusters”). To quantify coordinated lipid regulation within metabolic pathways, we computed edge modulation scores by first standardizing fold-change values to Z-scores across supertypes for each lipid, then calculating pairwise modulation strength as the product of absolute Z-scores between metabolically connected lipids. The analysis was restricted to connected components of the metabolic network containing ≥2 lipids, with zero-variance lipids excluded. Cluster-level modulation scores were obtained by averaging edge scores across all supertypes within each comodulation cluster.

### Sphingolipid signature

The sphingolipid signature (ceramides, hexosylceramides, and sphingomyelins) across 3x male, 3x non-pregnant female, and 3x pregnant female brains were computed by first z-score normalizing individual lipid species, then averaging by supertype and condition (male, non-pregnant female, female). This procedure is analogous to the one commonly used for gene expression analysis and implemented in Seurat^92^. After filtering for supertypes belonging to the core white matter class, the data were row-centered to remove supertype-specific baseline differences, allowing direct comparison of relative lipid abundances across biological conditions. The resulting per-supertype means for each condition were visualized using violin plots, connecting individual supertypes with line segments from male to non pregnant female, and from non pregnant female to pregnant female.

### Sample similarity analysis

For inter-sample similarity assessment, we computed *lipizone* supertype lipid centroids for each brain sample (3 male, 3 female, 3 pregnant female), concatenating these centroids to create a *lipizone*-aware lipidome representation per sample. After standardization, we analyzed these profiles using PCA and calculated sample-sample correlation matrices.

### Inter-individual variation analysis

We computed intraclass correlation coefficients (ICC) to assess inter-individual variation using a one-way random effects model, where ICC represents the proportion of total variance attributable to between-subject differences relative to within-subject variation. For each *lipizone* subclass and lipid, we calculated ICC values by comparing measurements across three male samples, estimating between-subject (σ^2^_b_) and within-subject (σ^2^_w_) variance components through analysis of variance decomposition. The ICC was computed as:

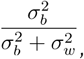

and averaged across subclasses, providing a metric of inter-individual variation for every lipid. We repeated the same procedure across three female samples, and averaged.

### Lipidome2Location model

The Lipidome2Location model is an out-of-the-box MLP (no hyperparameter tuning) to assess spatial predictability from lipidome profiles. The network uses 6 layers with ReLU activation, dropout (0.5, 0.3, 0.15, 0.1, 0.05), and batch normalization, trained for 50 epochs using MSE loss and Adam optimization with learning rate scheduling (ReduceLROnPlateau). The model was trained on 80% of right hemisphere sections of brain #1 with validation pixels, and tested on the remaining 20% and on the fully-held-out brain #2.

### Lipid brain atlas explorer

The Lipid Brain Atlas Explorer (LBAE; https://lbae-v2.epfl.ch/) is a Python-based interactive web application built using the Dash^93^ framework for visualizing and analyzing high-resolution mass spectrometry imaging data of lipids across the mouse brain. The LBAE modular architecture comprises data processing modules, visualization components, and interactive pages. The interface offers several analytical approaches: lipid selection for visualizing specific m/z peaks and annotated lipids; region analysis for comparing lipid abundances between neuroanatomical structures; *lipizone* visualization with ID cards inspection; lipid program analysis; and three-dimensional reconstruction of lipid distributions and *lipizones*. Interactive features include rostro-caudal slice navigation, region selection tools, volcano plots, and cross-sample comparative visualization. Real-time interaction with large-scale imaging datasets is enabled through strategic precomputation and caching mechanisms, making the platform and data accessible to researchers without specialized computational expertise.

## DATA AVAILABILITY

All data is granted public access. Raw data available on METASPACE at https://metaspace2020.org/project/mlba-2025 and processed data on Zenodo https://doi.org/10.5281/zenodo.15379499, https://doi.org/10.5281/zenodo.15379534, https://doi.org/10.5281/zenodo.15379565, in various formats including all metadata and in adata format for seamless integration with the scverse ecosystem^94^.

Interactive visualizations are available on our website at https://lbae-v2.epfl.ch/.

## CODE AVAILABILITY

EUCLID methods and tutorials are available as a PyPI package and a GitHub repository https://github.com/lamanno-epfl/EUCLID. The package is in active development and currently tested on a limited number of scenarios. The code to replicate the analysis of the paper and explore the dataset is available at https://github.com/lamanno-epfl/lipidbrainatlas. The code for lipiMap and the code to build the explorer are also available as separate GitHub repositories (https://github.com/lamanno-epfl/lipiMap, https://github.com/lamanno-epfl/lbae).

## ACKNOWLEDGEMENTS

L. F. B. is funded by a Boehringer Ingelheim Fonds PhD fellowship. G. D. A. is supported by the Swiss Cancer League (KFS-4999-02-2020), the EPFL institutional fund, the Kristian Gerhard Jebsen Foundation, and the Swiss National Science Foundation (SNSF) (310030_184926). G. L. M. is supported by SNSF (TMSGI3_218393, IZSEZ0_213427), SDSC grant agreement C22-07, the Kristian Gerhard Jebsen Foundation. Mice for LC-MS/MS were donated from Professor Johannes Gräff’s laboratory (license #3954). We thank Carmen Sandi, Carl Petersen, Amit Zeisel, Alessandro Furlan for constructive comments on the manuscript, Julijana Ivanisevic for the support on LC-MS, Gaël Rayot for the support with the website, and the members of the La Manno and D’Angelo Labs for their insightful comments.

## AUTHOR CONTRIBUTIONS

L. F. B. developed the idea, constructed the preprocessing pipelines, designed the algorithms and computational analyses, analyzed, visualized, and interpreted the data, designed the website, wrote the manuscript. H. H. S. developed the idea, constructed the preprocessing pipelines, and analyzed data. L. C. and L. H. A. A. conducted the MALDI-MSI experiments, performed image registration and contributed interpreting the data. F. V. designed the lipiMap algorithm and built the website. A. V. conducted the LC-MS experiment and interpreted the data. C. D. developed visualizations, contributed to registration and built the website. D. T. B. developed 3D interpolation and data visualizations. I. K. and E. Z. A. contributed to the establishment of the MALDI-MSI acquisition protocol on brain sections. A. N. and I. K. performed animal experimentation and brain harvesting. E. K. supervised clustering and 3D interpolation analysis. G. L. M. and G. D. A. developed the idea, secured funding, designed and supervised the entire project, analyzed and interpreted the data and wrote the manuscript. All coauthors read and approved the manuscript.

## ETHICS DECLARATIONS

None to declare.

## EXTENDED DATA FIGURES AND TABLES

**Supplementary Figure 1.**
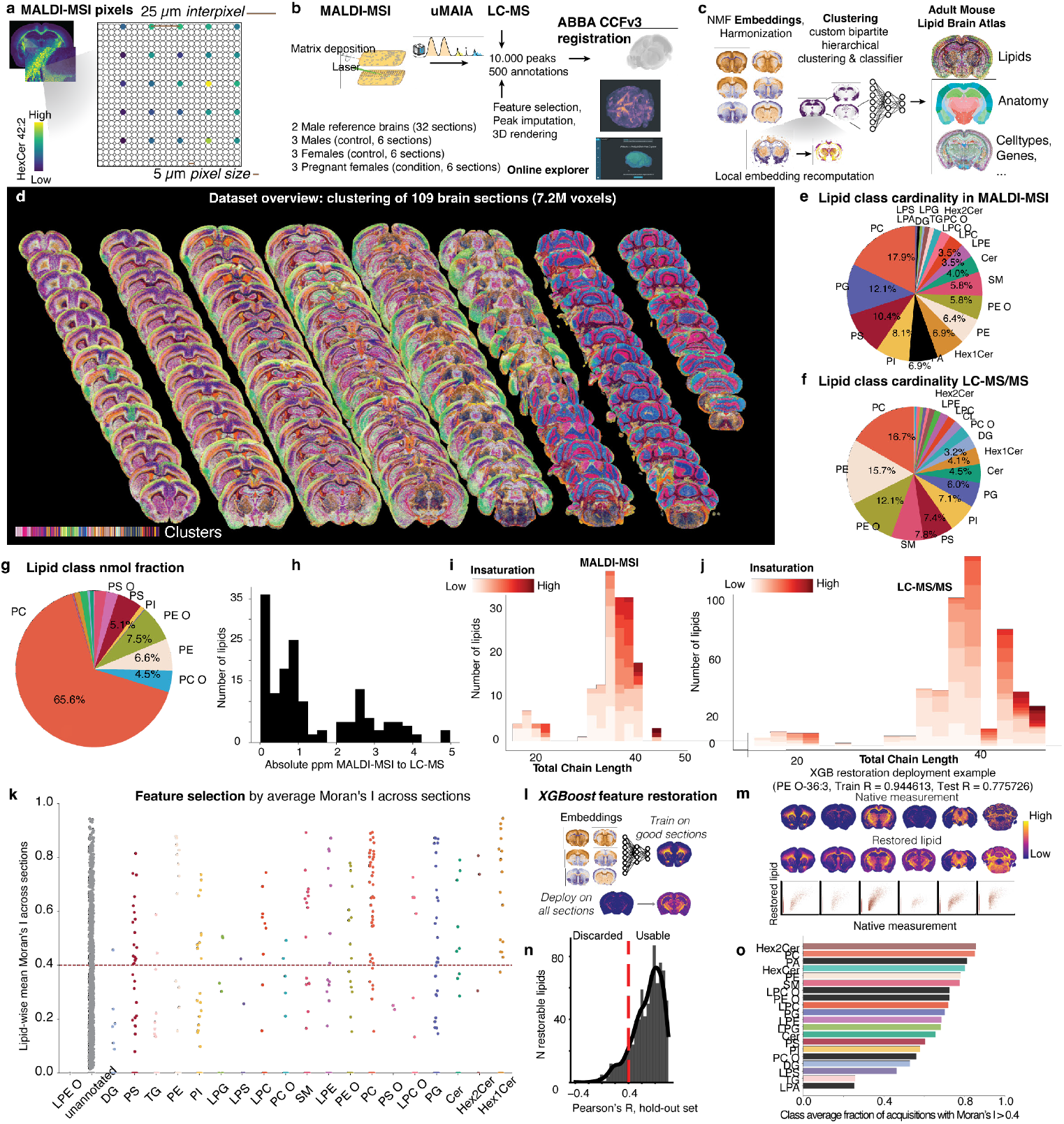
Study design, biochemical characterization of the lipids in this study, and feature restoration. (**a**) Spatial zoom-in representation of MALDI-MSI pixel sampling scheme, highlighting the pixel size and inter-pixel distance, colored by the abundance of the lipid HexCer 42:2. (**b**) Study design. (**c**) Computational processing overview. (**d**) Spatial plot for dataset overview, showing all brain sections colored by cluster (*lipizone*). (**e**) Pie chart for the proportions of lipid classes in terms of number of nonredundant lipids in this study (MALDI-MSI). (**f**) Pie chart for the proportions of lipid classes in this study in terms of number of nonredundant lipids for the paired whole-brain LC-MS. (**g**) Pie chart of nanomolar fractions for each lipid class in LC-MS. (**h**) Histogram for the absolute parts per million (ppm) distance between MALDI-MSI and LC-MS m/z peaks for the 172 nonredundant lipids used for the analysis in this study. (**i**) Nested histograms for carbon chain length and insaturation, for the lipids detected by MALDI-MSI in this study. (**j**) Nested histograms for carbon chain length and insaturation, in for the lipids detected by LC-MS in this study. (**k**) Strip plot of the Moran’s I scores for individual lipids in our MALDI-MSI dataset (average across sections), grouped by lipid class. (**l**) Schematic illustrating the XGBoost-based feature restoration strategy that we devised to recover lipid distribution estimates for sections with corrupted measurements. (**m**) Examples with spatial plots and reconstruction scatter plots measured vs restored of a lipid undergoing restoration (PE O-36:3). (**n**) Histogram of XGBoost performance on held-out data across lipids, evaluated by Pearson’s R. (**o**) Barplot for the average fraction of acquisitions with Moran’s I > 0.4, one bar per lipid class.

**Supplementary Figure 2.**
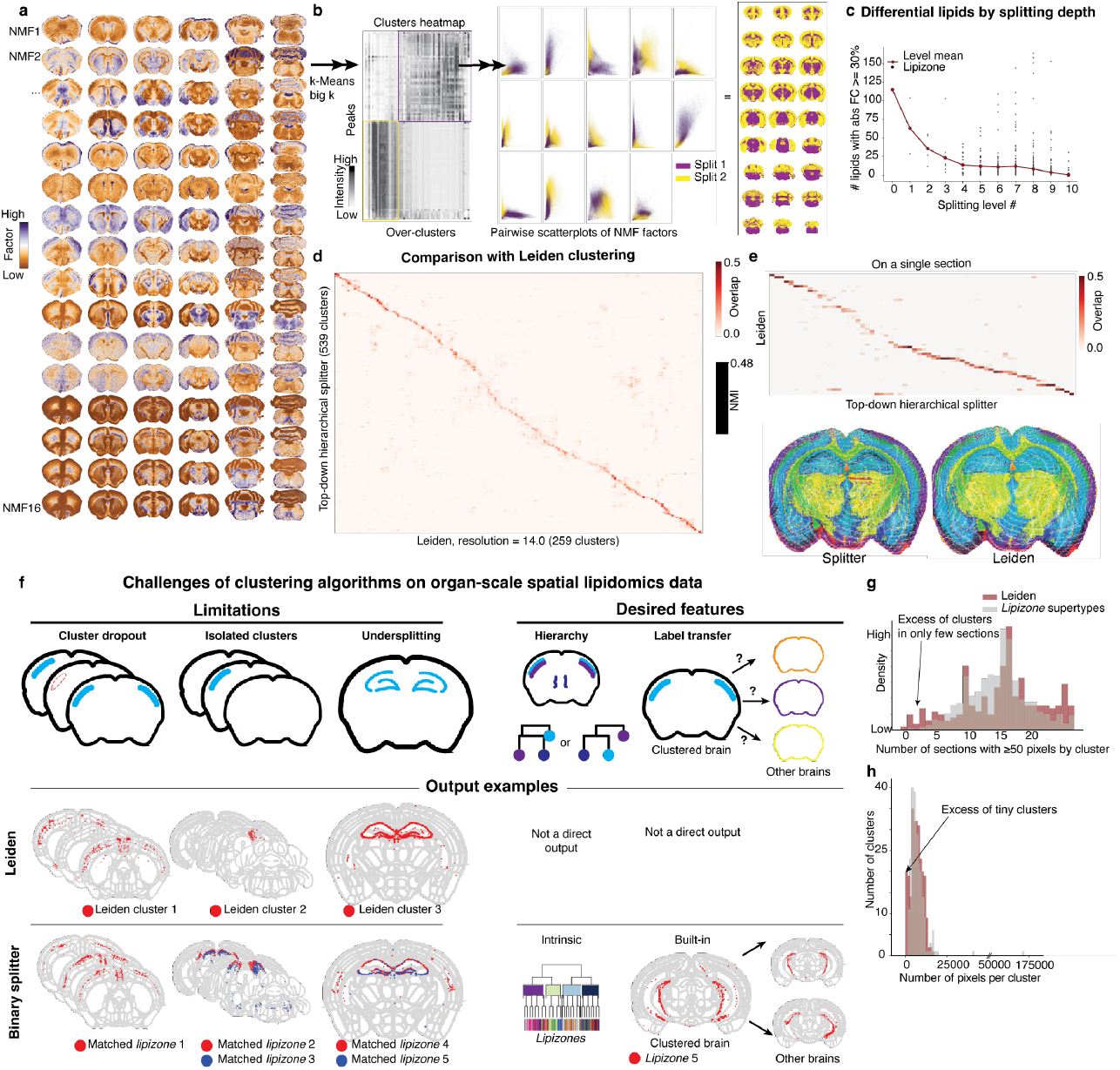
Embedding and clustering methods. (**a**) Spatial plot of the global, harmonized NMF embeddings for representative sections. (**b**) Left: hierarchical binary split procedure illustrated for the first split: heatmap of cluster centroids by m/z peaks sorted by backSPIN for cluster aggregation. The two resulting clusters are highlighted in violet and yellow. Center: scatterplot of the first binary split shown along NMF-pair scatters; right: the same clusters visualized spatially. (**c**) Strip plot of the number of lipids with absolute fold change over 30% by clustering splitting level, with overlaid mean trend. (**d**) Heatmap of sorted, normalized confusion matrix comparing the clustering method used in this study against Leiden clustering, evaluated across the entire brain. The inset shows the NMI score between the two clustering methods. (**e**) Heatmap of sorted, normalized confusion matrix comparing the clustering method used in this study against a Leiden clustering, on a single section, accompanied by color-matched spatial plots, where colors correspond to the clusters obtained from these two clustering strategies. (**f**) Vignette illustrating the challenges of clustering in organ-scale spatial lipidomics, and corresponding data for Leiden and the top-down custom binary splitter that we used. Leiden clustering was found to suffer from cluster dropout (i.e., clusters could skip some sections), isolation of clusters (implausible, given our dense section sampling scheme), and undersplitting (high residual within-cluster variability, that resulted in further splits using the binary splitter). (**g**) Frequency histogram for the number of sections a cluster reliably appears in for the binary splitter and Leiden clustering. (**h**) Frequency histogram of the cluster size variability for the clusters obtained with the binary splitter vs Leiden clustering.

**Supplementary Figure 3.**
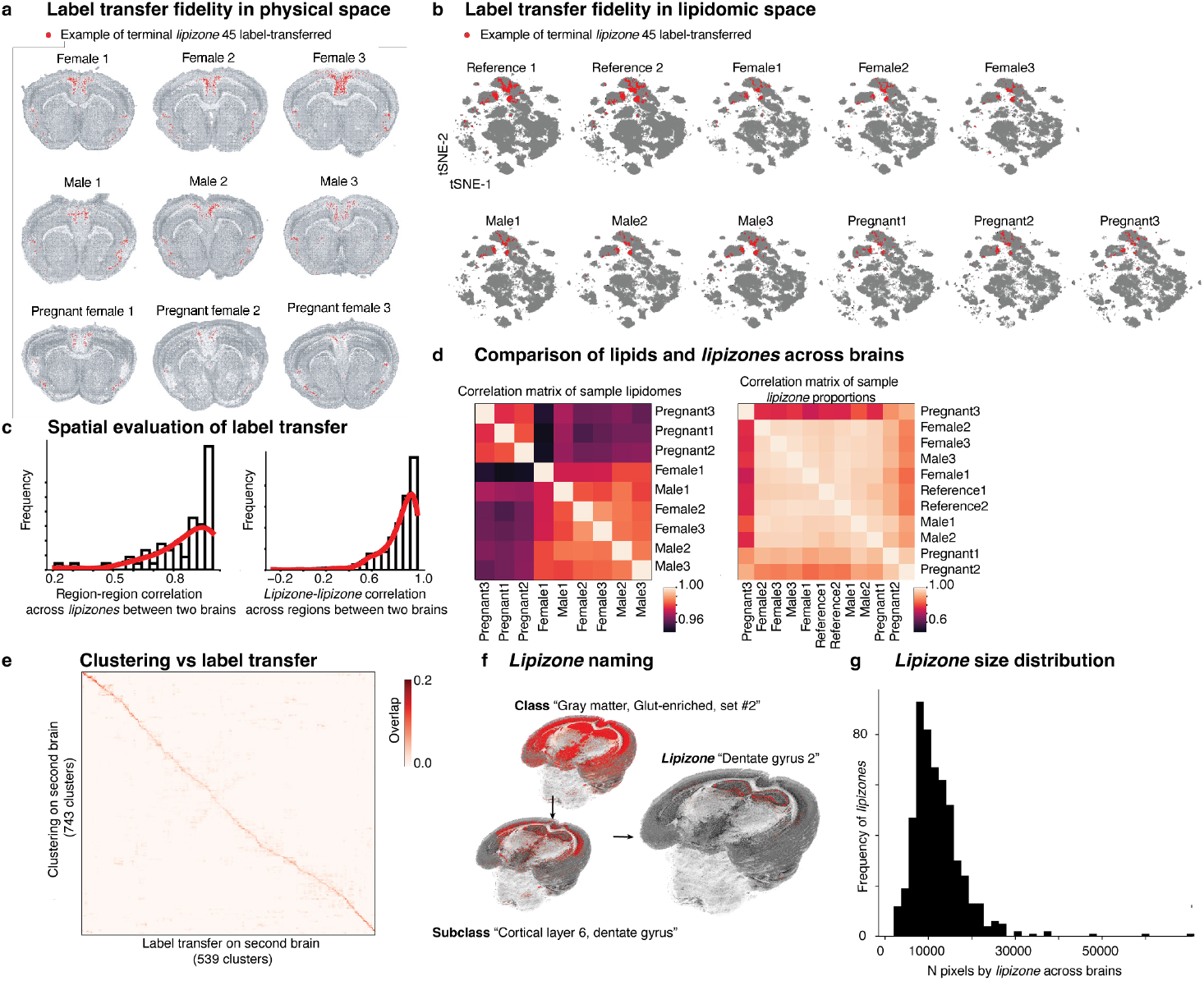
Label transfer, *lipizone* nomenclature, and inter-sample controls. (**a**) Spatial plot of a terminal example *lipizone* transferred across individuals showing spatial consistency. (**b**) t-SNEs computed from the lipidome, highlighting the same example *lipizone* as in (a), across individuals. (**c**) Frequency histograms of Pearson’s R for Allen region pairs between brain #1 and brain #2 in terms of *lipizone* enrichments and *lipizone* pairs in terms of Allen regions to explore fine-grained spatial consistency between the two brains after label transfer. (**d**) Heatmap of sample-sample correlation in terms of *lipizone* lipidomes and heatmap of sample-sample correlation in terms of *lipizone* proportions. (**e**) Heatmap of the normalized confusion matrix comparing reclustering and label transfer on brain #2. (**f**) Spatial plots for the *lipizone* hierarchy and naming convention at three hierarchical levels. (**g**) Histogram of the frequency distribution of the number of pixels per *lipizone*, pooling all the 11 brains.

**Supplementary Figure 4.**
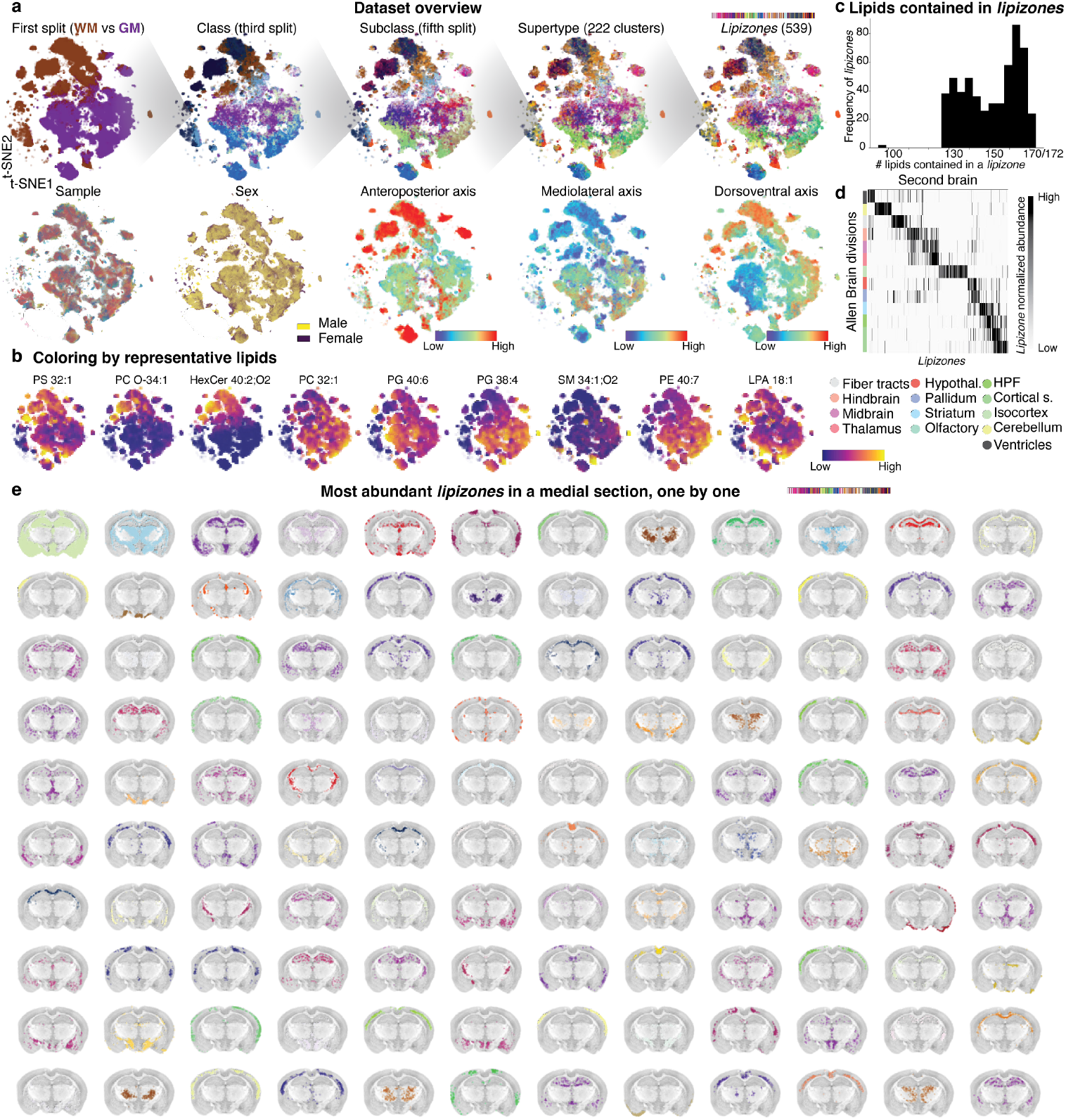
Dataset overview in feature space and physical space. (**a**) tSNEs colored by relevant clustering levels and metadata. (**b**) tSNEs colored by representative lipids. (**c**) Frequency histogram for the number of lipids contained in a *lipizone* across *lipizones*. (**d**) Heatmap replicating Figure 2e on brain #2, sorted by brain #1, representing *lipizone* normalized abundances by Allen Brain division. (**e**) Spatial plot of the most abundant *lipizones* in a single section, shown individually, color-coded.

**Supplementary Figure 5.**
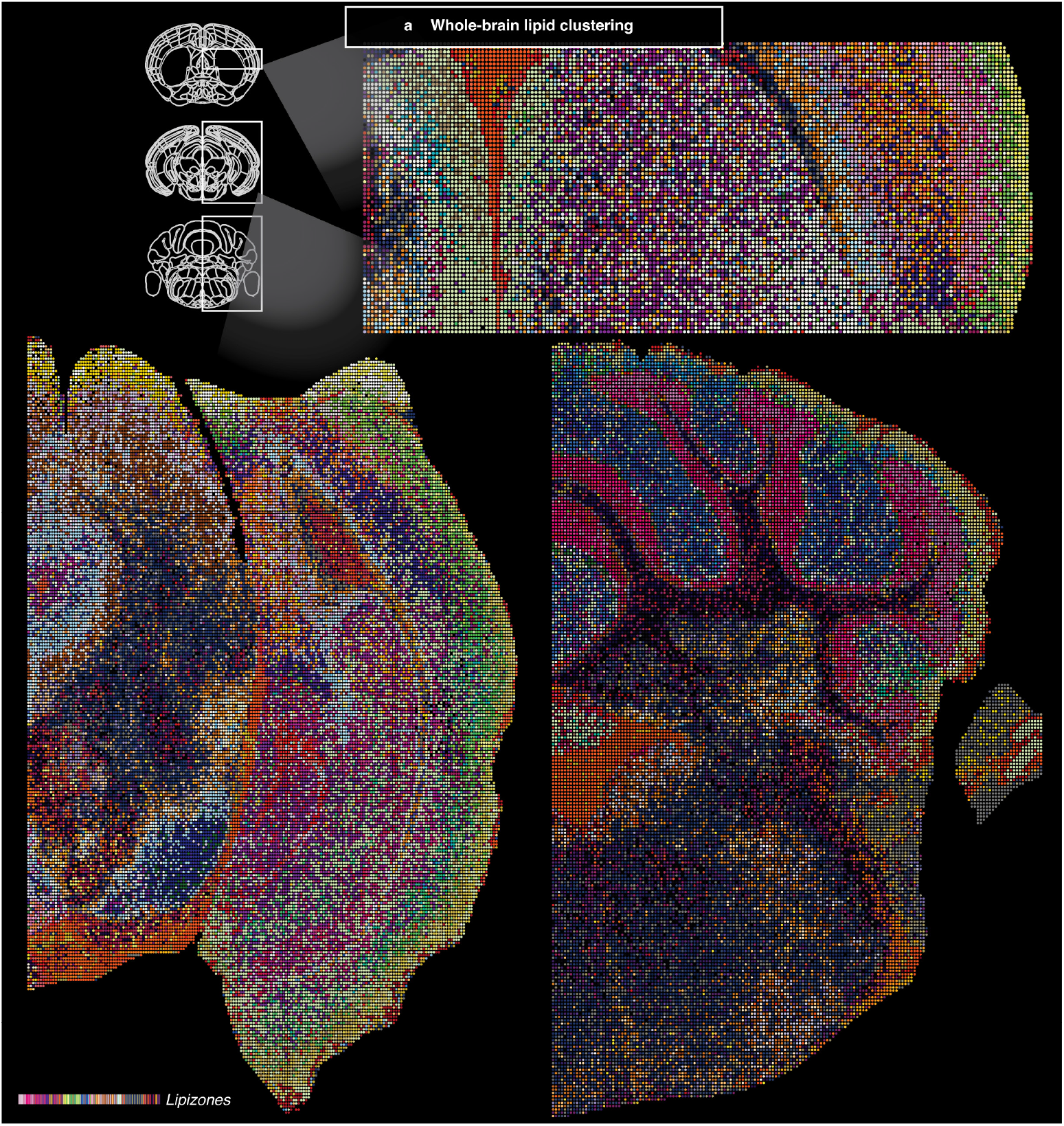
Mosaic plots for different brain regions. (**a**) Half-brain spatial plots spaced along the rostrocaudal axis, colored by *lipizone*, highlighting their spatial organization.

**Supplementary Figure 6.**
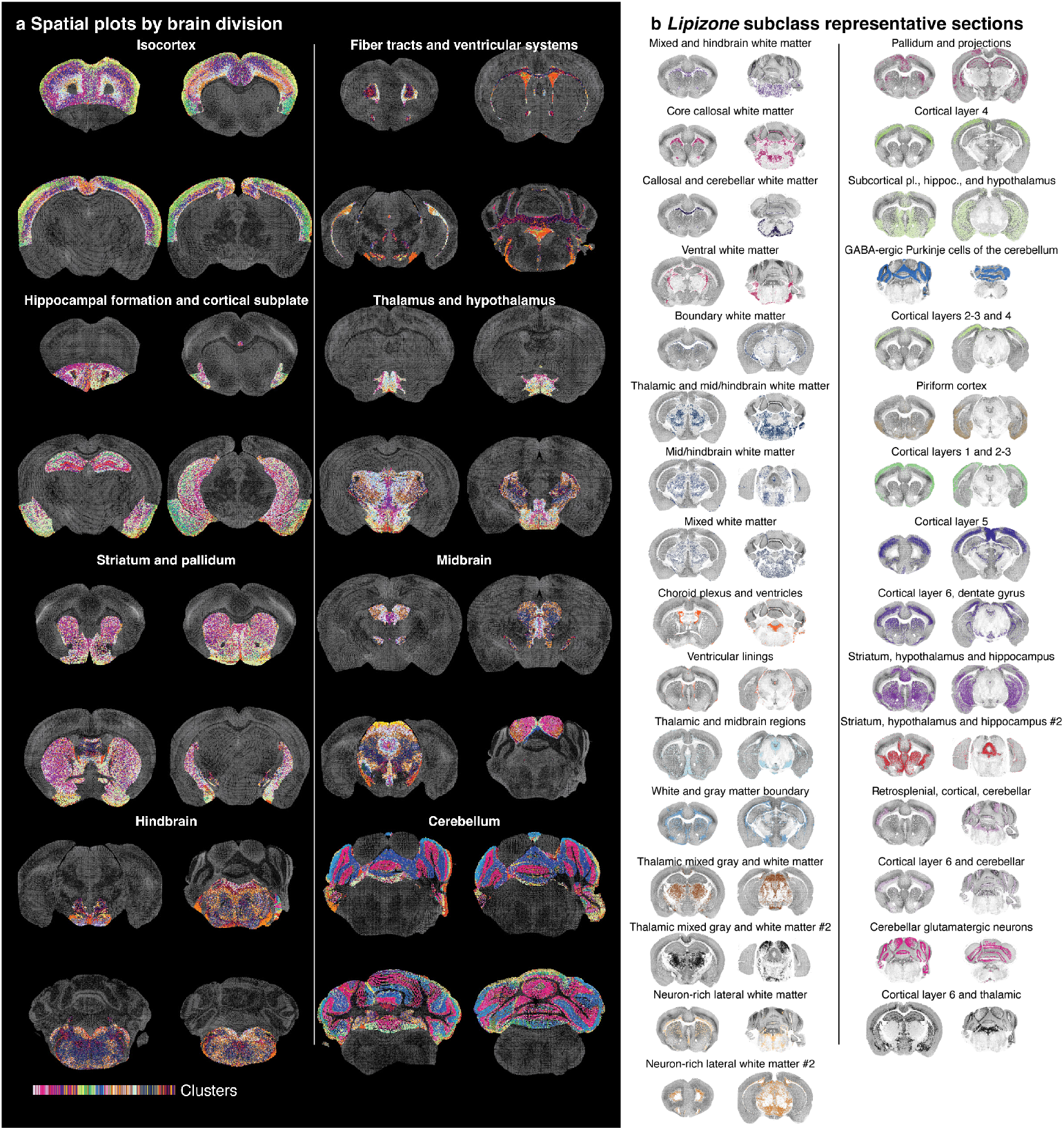
*Lipizones* spatial overviews by anatomical division and by lipidomic subclass. (**a**) Spatial plots of *lipizones* by brain division. (**b**) Spatial plot of the 31 lipid-derived subclasses of the brain for two representative sections.

**Supplementary Figure 7.**
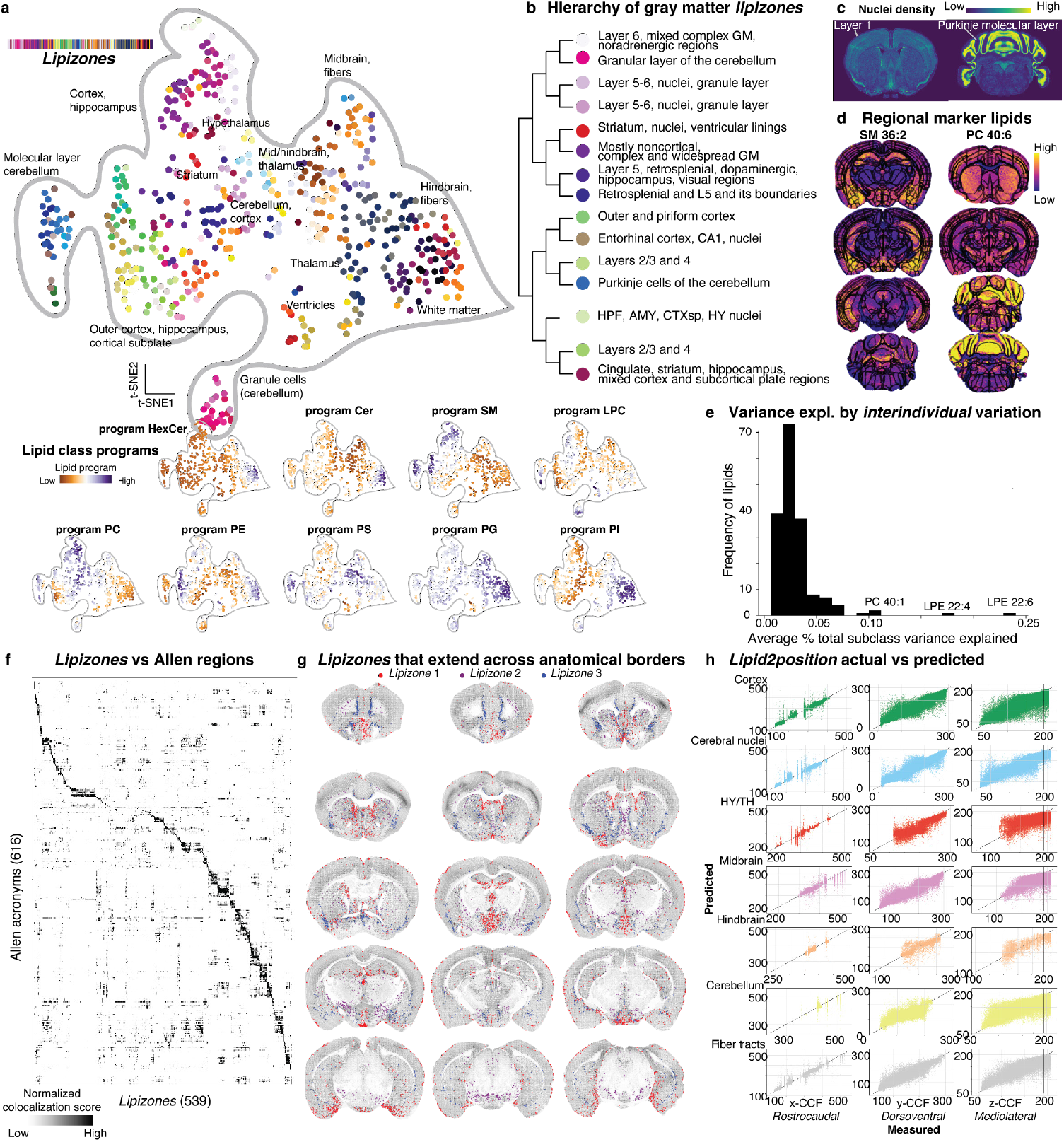
Organization of *lipizones* in terms of lipid classes and compared to brain anatomy. (**a**) t-SNEs of *lipizone* centroids, colored by *lipizones* and by the lipid programs representing lipid classes. (**b**) Dendrogram showing the early lipidomic splits in the gray matter-rich branch, manually labelled with anatomy-based names. (**c**) Representative Allen Brain nuclei density images showing low nuclei density in outer cortex and Purkinje molecular layer. (**d**) Spatial plots of two example regional marker lipids, SM 36:2 (cortical subplate and hippocampal formation) and PC 40:6 (cerebellum), for four representative sections along the anteroposterior axis each. A contour is overlaid, highlighting the CCF registration quality. (**e**) Frequency histogram for the average percentage of total subclass variance explained by inter-individual variation, across lipids. (**f**) Heatmap of the colocalization score (reciprocal enrichment) between *lipizones* and Allen Brain regions. (**g**) Spatial plot example of 3 *lipizones* with geometrical but nontrivial distributions crossing conventional anatomical boundaries, shown together across 15 representative sections, using 3 colors. (**h**) Pairwise scatterplots for the ground truth and lipidome-reconstructed 3D spatial coordinates by the “*Lipid2position*” model (1 column per brain axis); one point is a brain pixel; subplots are organized and colored by Allen Brain division.

**Supplementary Figure 8.**
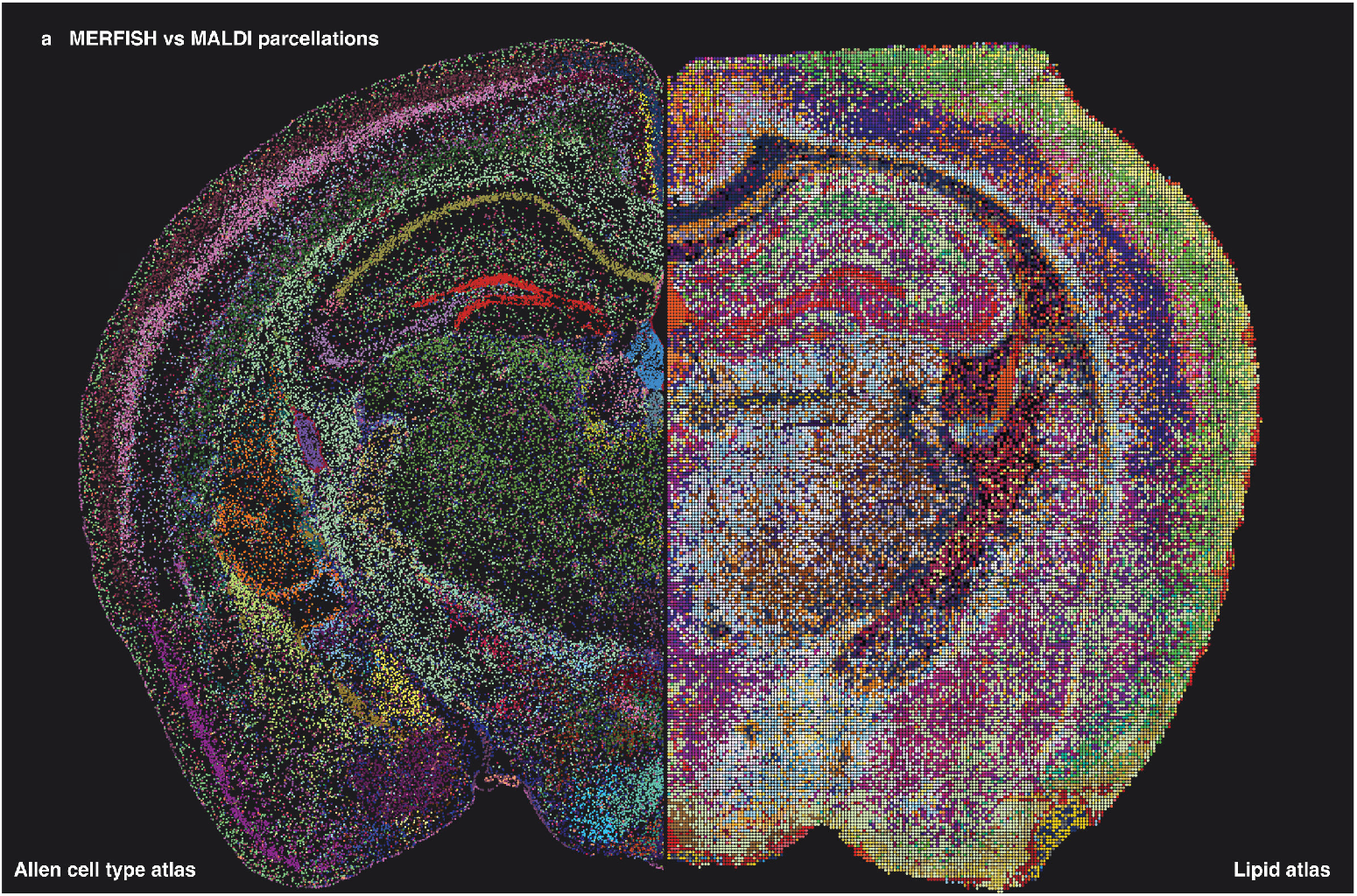
Visual comparison of the Allen Brain cell type atlas and the lipid brain atlas. (**a**) Half-brain spatial transcriptomics parcellation and half-brain spatial lipidomics parcellation, colored by subclass-level cell types, and *lipizones*, respectively.

**Supplementary Figure 9.**
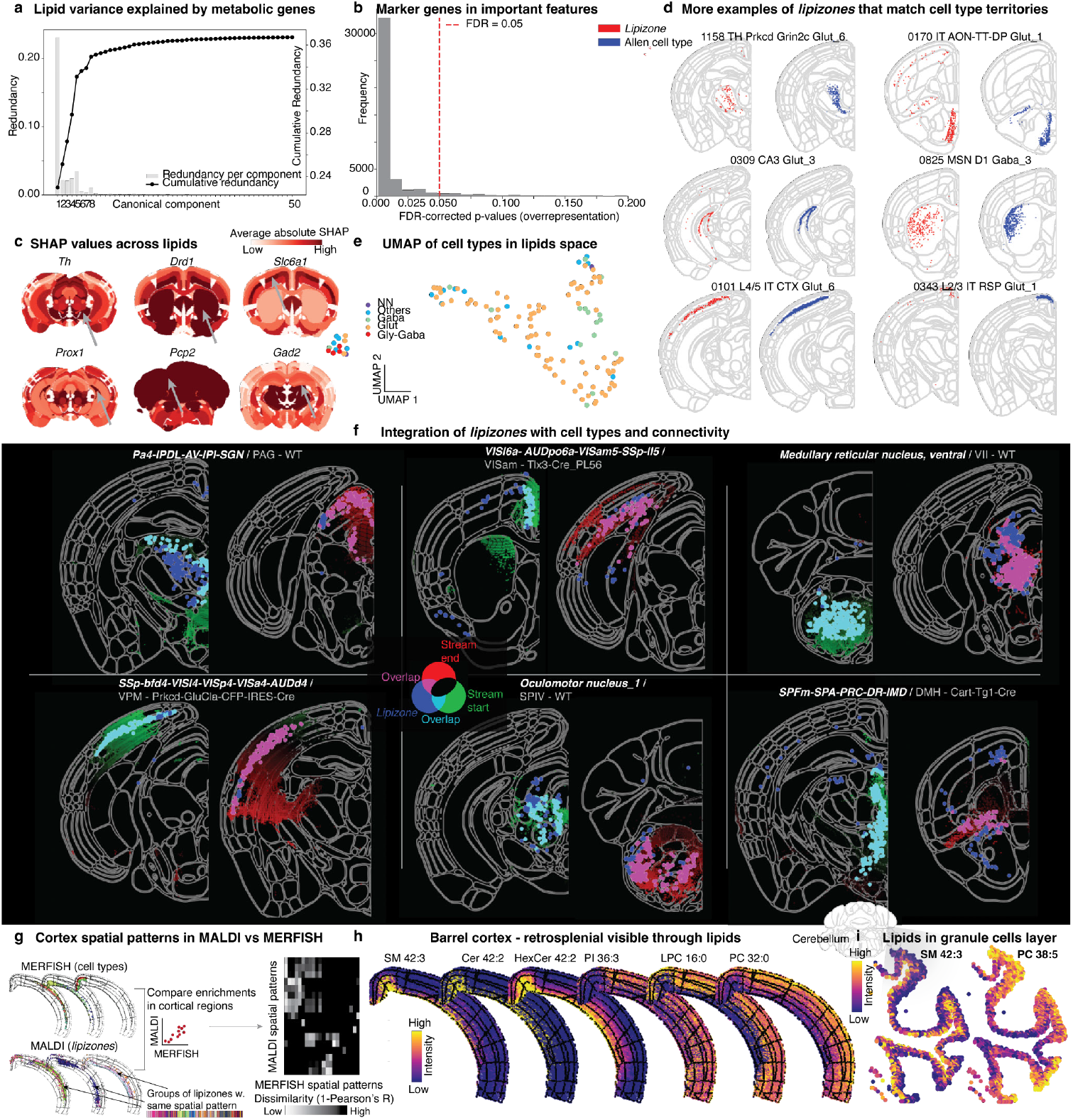
Multimodal analysis and organizational principles of *lipizone* distribution. (**a**) Redundancy-scree plot showing per-component and cumulative lipid variance explained by metabolic gene-derived canonical variates (canonical component analysis using the Allen 8460-genes imputed transcriptome and the lipid brain atlas, matching voxels in physical space). (**b**) Frequency histogram of BH-corrected FDRs for cell type marker gene overrepresentation in the top 20 important features for the XGBoost gene to lipid regression, across all lipids and anatomical regions, showing that, for the large majority of lipid-region pairs, cell type markers are largely enriched in the top important features set. (**c**) Spatial plot examples of SHAP feature importance maps for the gene to lipid XGBoost regression, averaged across all lipids. Shown are selected well-known cell type regional marker genes. (**d**) Example spatial plots of *lipizones* and the cell type territories they overlap with. (**e**) UMAP of cell type centroids computed from their estimated lipidome. (**f**) 2D visualizations of six further representative *lipizones* that likely include cell bodies (cyan) and their distal terminals (magenta). The associated connectomic streams are represented in background, colored from green to red, passing through black. The pixels of the *lipizones* that do not overlap with the connectomic streams are colored in blue. The gray text in the titles, after the white *lipizone* names, contains the connectome experiment injecton site and mouse line. (**g**) Left, schematic of the experiment comparing the unique spatial patterns delineated by MERFISH-derived cell types and MALDI-derived *lipizones*. We assessed the enrichment of each cell type and lipizone across cortical Allen regions, clustered cell types and *lipizones* based on these regional enrichments to detect unique spatial patterns, and compared these clusters between MERFISH and MALDI. Right, corresponding heatmap quantifying the pairwise dissimilarity of MERFISH cell type spatial patterns and MALDI *lipizone* spatial patterns in the cortex, evaluated as 1 - Pearson’s R calculated on the regional enrichments. (**h**) Spatial plots restricted to half section cortex for representative lipids highlighting the barrel and retrosplenial common lipidome captured by *lipizones*. (**i**) Zoom-in spatial plots for the cerebellar granule cells layer, detected as two major layered *lipizones*, colored by two lipids that were differential between the inner and outer layer *lipizones*.

**Supplementary Figure 10.**
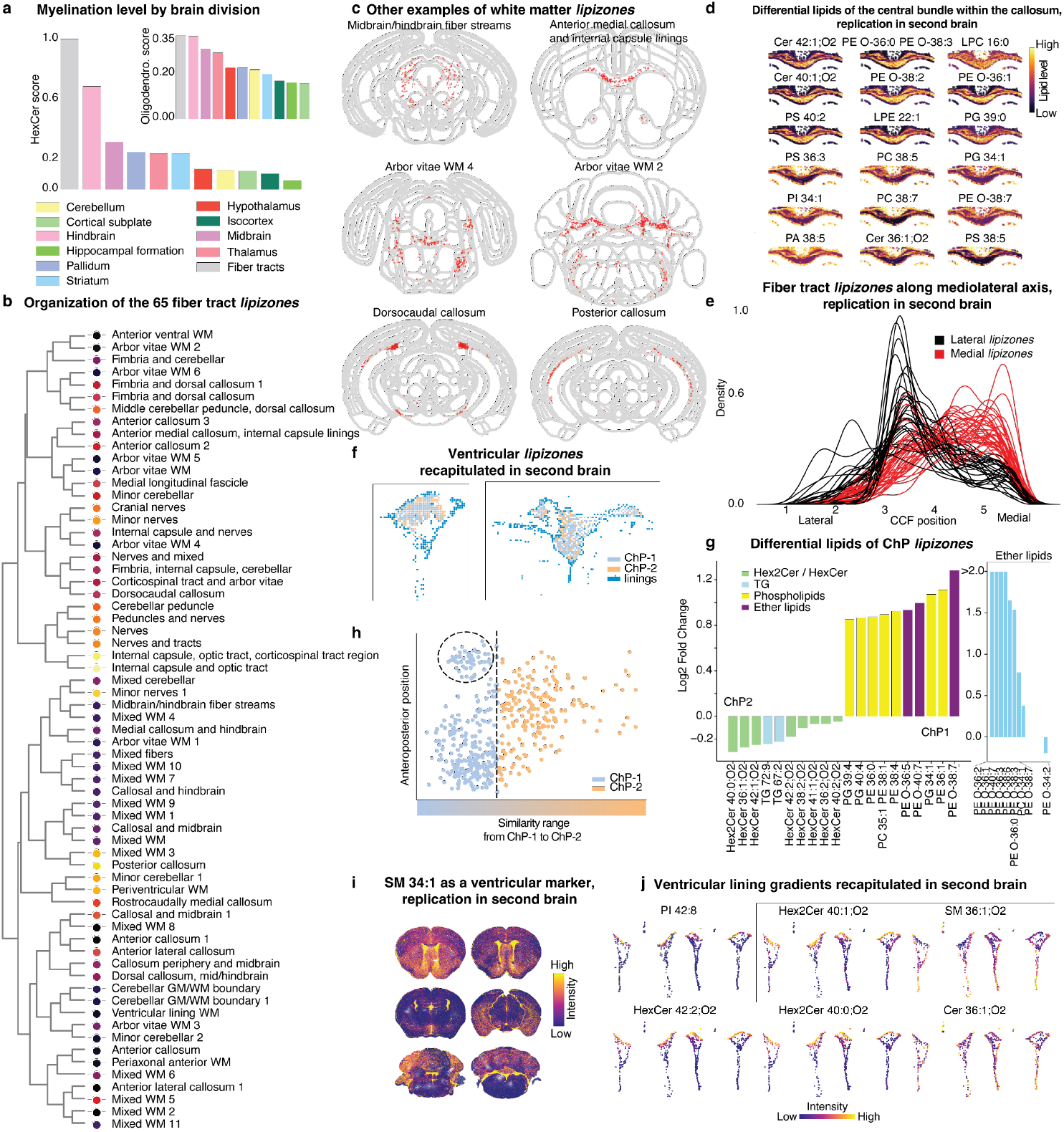
Lipidomic features of the white matter and the ventricular systems. (**a**) Barplot of myelination and oligodendrocyte scores for each Allen division, sorted and colored by Allen division color. (**b**) Annotated *lipizones* tree for the 65 core white matter-related *lipizones*. (**c**) Spatial plots of example regionalized white matter-related *lipizones*, in addition to those in Figure 4. (**d**) Zoom-in spatial plots for replication of the axonal bundle marker lipids on brain #2. (**e**) Kernel density estimation for replication on brain #2 of the mediolateral pattern of the 65 core white matter-related *lipizones* in the anterior callosum. (**f**) Spatial plot replication of the ChP1, ChP2, and lining *lipizones*, on brain #2. (**g**) Left, barplot of ChP1 and ChP2 marker lipids; right, lightblue-colored barplot showing ether lipid imbalance. (**h**) Scatterplot of the similarity range vs anteroposterior position for whole-brain *lipizones* against ChP1-2, showing the cerebellar *lipizones* omitted from the analysis of Figure 5e. (**i**) Spatial plot replication of SM 34:1 as a ventricular marker, on brain #2. (**j**) Zoom-in spatial plot replication of the dorsoventral lipid gradients along the ventricular walls, on brain #2.

**Supplementary Figure 11.**
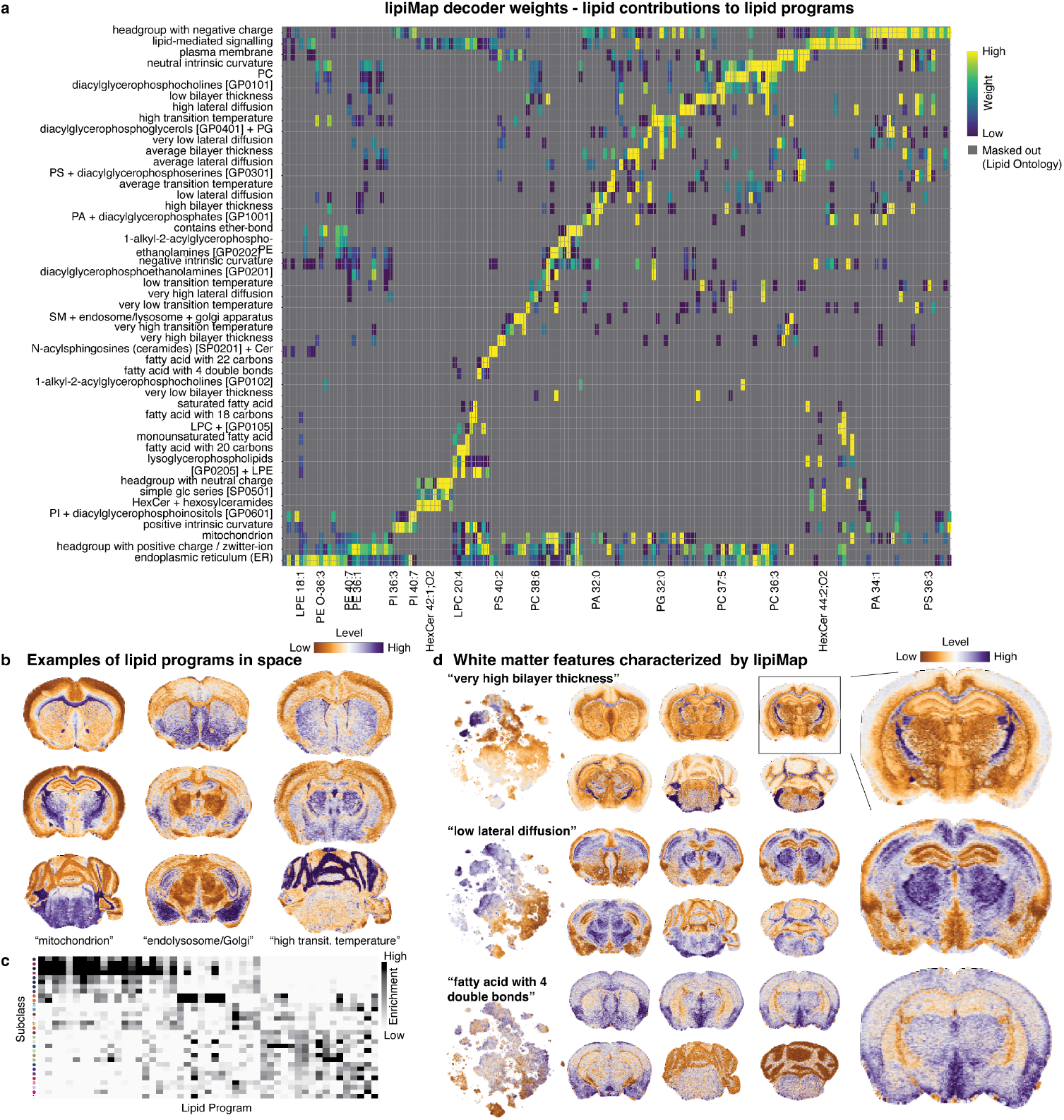
A biochemically-constrained variational autoencoder to disentangle lipid programs. (**a**) Heatmap of the decoder weights of the variational autoencoder, for each lipid program and lipid pair, gray where a lipid does not belong to a program; color intensity otherwise represents the weight of a lipid program to reconstruct the abundance of a given lipid. Columns are individual lipids; only few lipids are labelled for readability. (**b**) Spatial plots for example lipid programs, representative sections. (**c**) Heatmap of lipid program enrichments by *lipizone* subclass. (**d**) tSNEs and spatial plots of representative sections, with zoom-ins, colored by lipid programs that characterize the white matter and its heterogeneity.

**Supplementary Figure 12.**
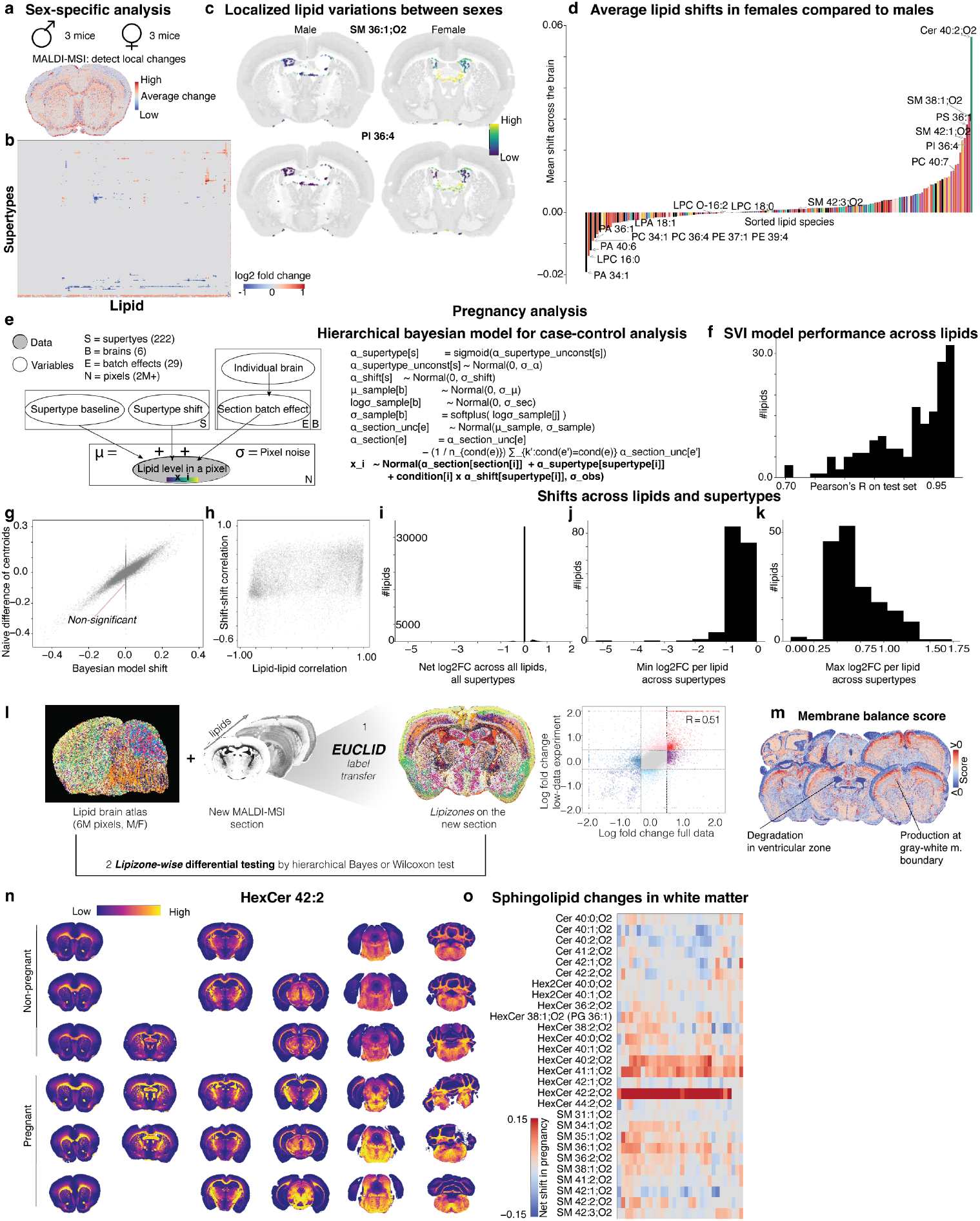
Localized variation between sexes and in pregnancy discovered with Bayesian modeling. (**a**) Spatial plot for the overall change in lipidome between sexes as obtained by summing the net female-male differences across lipids, shown for an anterior section. (**b**) Heatmap of log2FCs for all lipids in all supertypes between males and females. (**c**) Spatial plots of localized lipid differences between males and females, two example lipids in a ventricular *lipizone*. (**d**) Barplot showing the whole-brain average lipid change between males and females for all the lipids in this study, colored by lipid class. (**e**) Schematic of the hierarchical Bayesian model used to detect supertype-specific changes in pregnancy and between sexes, while accounting for interindividual variation and batch effects. (**f**) Histogram of Pearson’s R on *lipizone* centroids for the Bayesian model SVI reconstruction performance across lipids in pregnancy, evaluated as Pearson’s R between the ground truth lipid abundance and its posterior average. (**g**) Scatterplot of the Bayesian model shifts and the naïve empirical differences of average in pregnant mouse brains and average in non pregnant mouse brains, across all lipids and supertypes. (**h**) Scatterplot of lipid-lipid correlation versus the correlations of the same lipid pairs calculated from their changes in response to pregnancy, showing that covariation in response to the pregnancy perturbation does not trivially reflect the baseline lipid covariation patterns in the brain. (**i-k**) Frequency histograms for the average, minimum, and maximum fold-change per lipid across supertypes. (**l**) Schematic of EUCLID algorithms use in low-data settings (label transfer of *lipizones*, Bayesian model), and scatterplot comparing the log2FCs obtained when using the whole dataset with the Bayesian model versus the log2FCs estimated empirically from an individual section of a pregnant mouse brain using the atlas. (**m**) Spatial plot of the membrane balance score for representative sections, which quantifies the supertype-resolved overall lipid variation in pregnancy, by summing the net shifts inferred via Bayesian modeling across all lipids, charting the relative prevalence of biosynthesis or turnover. (**n**) Spatial plot showing all the sections, in non-pregnant and pregnant mouse brains, colored by the white matter marker HexCer 42:2. (**o**) Heatmap of empirical sphingolipid net shifts in pregnancy across core white matter supertypes. Rows are sorted by class, chain length, and insaturation.

